# Controlling mechanism of the Scc2-cohesin interaction to restrict peri-centromeric DNA loop expansion and facilitate mitotic chromosome segregation

**DOI:** 10.1101/2024.04.05.588214

**Authors:** Sao Anh Nguyen, Toyonori Sakata, Katsuhiko Shirahige, Takashi Sutani

**Affiliations:** Institute for Quantitative Biosciences, The University of Tokyo 1-1-1 Yayoi, Bunkyo-Ku, Tokyo, 113-0032, Japan; Department of Cell and Molecular Biology, Karolinska Institutet, Sweden 171 77, Stockholm, Sweden; Department of Biosciences and Nutrition, Karolinska Institutet, Sweden 171 77, Stockholm, Sweden

**Keywords:** Cohesin, DNA loop, loop extrusion, budding yeast, Micro-C, centromere, chromosome segregation, NIPBL, WAPL, Eco1

## Abstract

Cohesin exhibits DNA loop extrusion activity when bound to an ATPase activator Scc2 (NIPBL in humans), thereby organizing higher-order chromosome folding. In budding yeast, the majority of the chromosome-bound cohesins lack association with Scc2. It remains unknown how the interaction between Scc2 and cohesin is regulated on the chromosome and what physiological consequences malfunction in this regulatory mechanism causes. Here, we show that simultaneous deletion of Wpl1 and Eco1, two of the known cohesin regulators, resulted in Scc2 co-localization with cohesin around the centromeres of metaphase chromosomes. In these cells, the pericentromeric DNA loops were enlarged to connect the centromere to the cohesin/Scc2 co-bound sites at distances of up to a few hundred kb, indicating highly active loop extrusion by the Scc2-associated cohesin. Furthermore, we demonstrated that Δ*wpl1* Δ*eco1* cells exhibited a delay in the progression of mitotic chromosome segregation, a phenotype dependent on the presence of Scc2 in metaphase. These findings suggest that Wpl1 and Eco1 cooperatively regulate Scc2-cohesin interaction on chromosomes, restrict the size of the pericentromeric DNA loops, and facilitate mitotic chromosome segregation.

## Introduction

Recent developments in genome-wide chromosomal conformation capture techniques, such as the Hi-C method, have revealed the principle of higher-order chromosome folding. The formation of DNA loops and sub-Mb-scale self-interacting regions (i.e., regions within which DNA interacts with each other at high frequency) have been found in various species, including yeasts and mammals, and are thought to be a fundamental feature of chromosome structure (Davidson & Peters, 2021; Fudenberg et al., 2017; Yatskevich et al., 2019). A prominent example of the self-interacting domains is the TAD (topologically associated domain) found in mammalian nuclei (Lieberman-Aiden et al., 2009).

Cohesin is one of the structural maintenance of chromosomes (SMC) protein complexes and plays a significant role in the formation of DNA loops and TAD structures (Davidson & Peters, 2021; Hoencamp & Rowland, 2023; Yatskevich et al., 2019). It is composed of four subunits, Smc1, Smc3, Scc1/Rad21, and Scc3 (SA1 or SA2 in humans) and exhibits an overall ring-like configuration. Recently, cohesin was shown to introduce loop structures to DNA in the presence of ATP and Scc2-Scc4 dimer (NIPBL-MAU2), an activator of cohesin ATPase (Davidson et al., 2019; Davidson & Peters, 2021; Kim et al., 2019). Scc2 directly interacts with cohesin at multiple interfaces. In the loop extrusion reaction, cohesin first forms a small loop at the DNA site it is bound to, then gradually increases the size of the loop by extruding DNA through its ring. Loop extrusion by cohesin can be impeded by various roadblocks on DNA. A typical example is the CTCF protein, whose binding sites are enriched at TAD boundaries (Davidson et al., 2023; Dixon et al., 2012; Nora et al., 2017; Rao et al., 2014; Zuin et al., 2014). The structural features of the TAD observed by Hi-C are well explained by the model that the cohesin-mediated expansion of DNA loops starting at any point in the genome is restricted by the CTCF binding sites (Fudenberg et al., 2016). In addition to TAD formation, cohesin-mediated loop extrusion is reportedly important for transcription control, DNA repair, and V(D)J recombination processes in mammals (Davidson & Peters, 2021; Hoencamp & Rowland, 2023; Peters, 2021).

Cohesin activity is regulated by various factors other than Scc2. Pds5 binds to cohesin in a mutually exclusive manner with Scc2, and the binding of Pds5 makes cohesin ATPase inactive (Kikuchi et al., 2016; Petela et al., 2018). Wpl1 (WAPL) interacts with cohesin via Pds5 and induces cohesin dissociation from chromosomes (Gandhi et al., 2006; Kueng et al., 2006; Rowland et al., 2009; Shintomi & Hirano, 2009; Sutani et al., 2009). Before its function in loop extrusion was discovered, cohesin was known for its canonical role in sister chromatid cohesion (K. Nasmyth & Haering, 2009). The establishment of sister chromatid cohesion requires acetylation of the Smc3 subunit by Eco1 (ESCO1 or ESCO2) in a manner coupled with DNA replication (Ben-Shahar et al., 2008; Lengronne et al., 2006; Rowland et al., 2009; Ünal et al., 2008). The acetylated cohesin reveals weakened interaction with Scc2 (Kaushik et al., 2023; Shi et al., 2020). The size of the cohesin-mediated loops is expected to vary depending on the affinity of cohesin for Scc2 and the retention time of cohesin on DNA. Indeed, in human cells lacking PDS5 or WAPL, the size of cohesin-mediated loops is larger, and loops are more easily formed to overcome barriers caused by CTCF (Gassler et al., 2017; Haarhuis et al., 2017; Wutz et al., 2017).

Cohesin-dependent loops and self-interacting regions are also found in chromosomes of budding yeast *Saccharomyces cerevisiae* (Costantino et al., 2020; Dauban et al., 2020; Hsieh et al., 2015; Lazar-Stefanita et al., 2017). CTCF is not present in this organism, and the boundaries of the self-interacting regions are defined by other means (Jeppsson et al., 2022). Cohesin loop expansion is negatively regulated by Pds5, Wpl1, and Eco1 in yeast (Bastié et al., 2022; Dauban et al., 2020), indicating the regulation of DNA loop enlargement is highly conserved between yeast and mammals. In wild-type (wt) mitotic cells whose centromeres are dissociated from microtubules, another prominent chromosome structure has been reported; the flanking regions adjacent to the centromere are linked with nearby cohesin binding sites to form so-called pericentromeric loops (Paldi et al., 2020). These pericentromeric DNA loops are generated presumably by cohesin loaded at the core centromeres that are unattached to the mitotic spindle. The expansion of the loops is confined by the genes that are 10–30 kb away from and transcribed towards the centromeres, and this confinement ensures the maintenance of sister chromatid cohesion in regions distal to the centromeres. Reorientation of the border-defining genes resulted in the loss of cohesion in wider genome region and impairs the establishment of chromosome biorientation under the spindle tension (Paldi et al., 2020).

A remarked difference in the behavior of budding yeast cohesin from that of mammalian one is that the vast majority of cohesin on the genome is not bound to Scc2/Nipbl (Hu et al., 2011; Kagey et al., 2010) and presumably exhibits only limited ATPase activity. This may be related to the relatively small size of the DNA loops in this organism (Hsieh et al., 2015). It remains to be understood how the Scc2-cohesin interaction is controlled on budding yeast chromosomes and what physiological consequences malfunction in the control mechanism causes. Here, we revealed that simultaneous deletion of Wpl1 and Eco1 resulted in co-localization of Scc2 with cohesin specifically on mitotic chromosomes without spindle tension. The same Δ*wpl1* Δ*eco1* cells exhibited extensively enlarged pericentromeric DNA loops, whose anchor sites corresponded to the co-localization sites of Scc2 and cohesin. The formation of the enlarged loops depended on the presence of Scc2 in mitosis, and Wpl1 and Eco1 synergistically contributed to the loop size restriction. Finally, we found that Δ*wpl1* Δ*eco1* cells showed delayed progression of chromosome segregation and that this phenotype was rescued by acute depletion of Scc2 in mitosis. Collectively, the data demonstrates that Wpl1 and Eco1 cooperatively control Scc2-cohesin interaction, restrict pericentromeric DNA loop expansion, and facilitate unperturbed chromosome segregation.

## Results

### Simultaneous depletion of Wpl1 and Eco1 promoted Scc2 co-localization at the cohesin binding sites

We explored how the genome-wide localization of Scc2, the cohesin ATPase activator, was affected by mutations in the cohesin regulators by spike-in ChIP-seq. First, we analyzed the localization in the cells arrested in metaphase by benomyl treatment. Proper arrest was verified by flow cytometry analysis (Figure S1). Chromosomal binding of Scc2 was displayed using normalized fold-enrichment (nFE), or ChIP/input ratio after normalization by spike-in control. In wt metaphase cells, centromeres and the surrounding regions are the major Scc2 binding sites (Figure 1A). On chromosome arms, Scc2 binding sites were coincident with those of Rpo21, a subunit of RNA polymerase II, most of which were not occupied by a cohesin subunit Scc1 (Figures 1A and S2A, B). These binding sites are identical to those already reported and assumed to correspond to the cohesin loading sites (Hu et al., 2011; Lengronne et al., 2004). After the loading reaction, cohesin presumably loses the binding to Scc2 and translocases to its chromosomal binding sites between convergently oriented gene pairs by passive diffusion (Lengronne et al., 2004). We found that double deletion of *WPL1* and *ECO1* (Δ*wpl1* Δ*eco1*) resulted in novel, prominent peaks of Scc2 along chromosome arms in metaphase (Figure 1A). Note that *ECO1* is an essential gene, but Δ*eco1* becomes viable by simultaneous deletion of *WPL1* gene (Sutani et al., 2009). The number of peaks in the genome was 225, 42% more than that for wt (Figure S2B). Notably, the majority (67%) of Scc2 peaks in Δ*wpl1* Δ*eco1* overlapped with Scc1’s peaks (Figure S2B). Co-localization of Scc1 and Scc2 in Δ*wpl1* Δ*eco1* was confirmed by correlation analysis; Spearman’s correlation coefficient increased from 0.16 in wt to 0.49 in Δ*wpl1* Δ*eco1* (Figure S2C). Spike-in ChIP-seq of Scc1 and Scc2 showed a high correlation between two biological replications, both for wt and Δ*wpl1* Δ*eco1* (Figure S3A).

**Figure 1.**
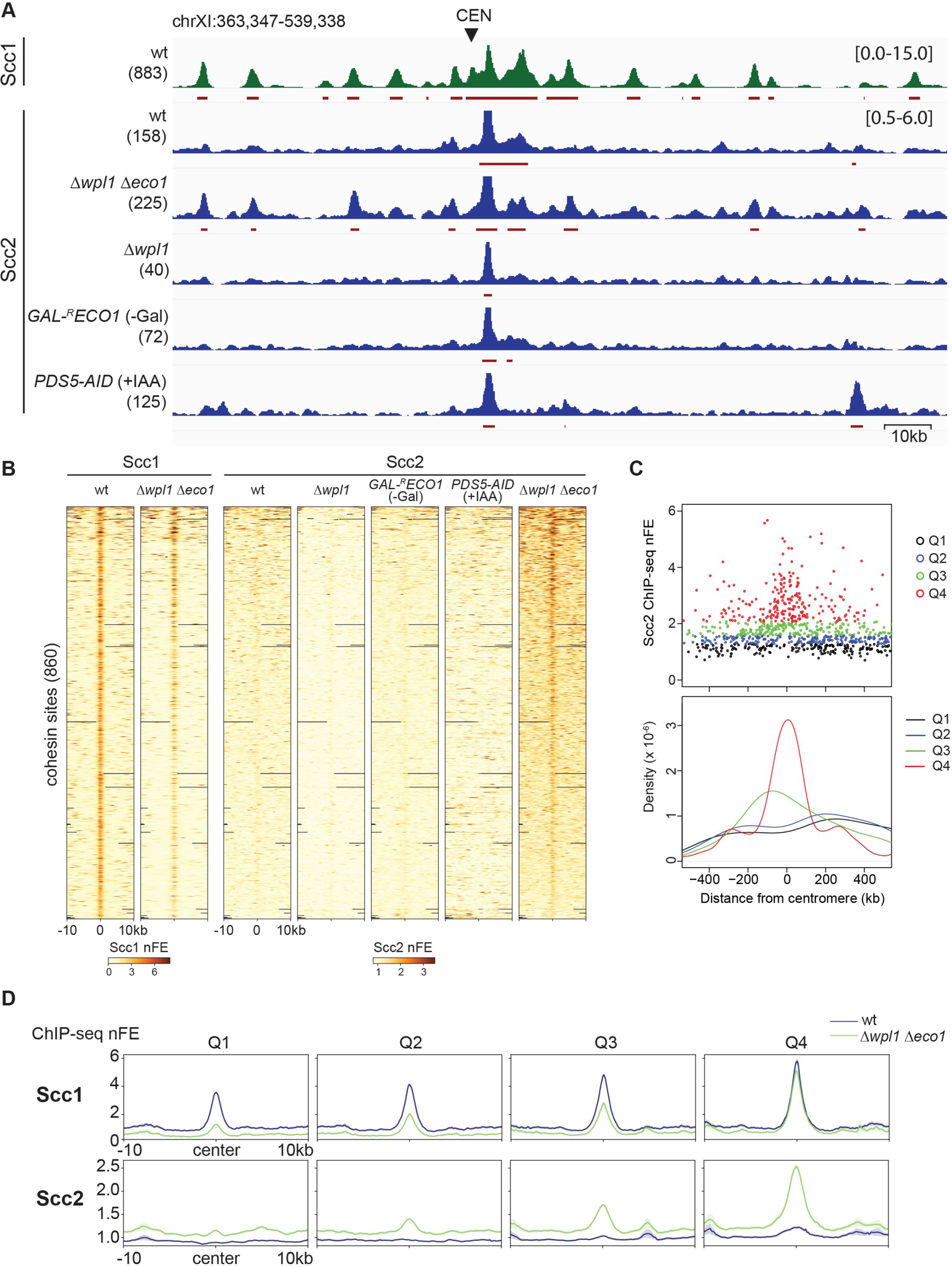
Simultaneous depletion of Wpl1 and Eco1 promotes Scc2 co-localization at cohesin binding sites. **(A)** Calibrated ChIP-seq profiles of Scc1-PK in wild-type (wt, strain SN75) and Scc2-PK in wt (SN40), Δ*wpl1* Δ*eco1* (SN54), *Δwpl1* (SN53), *GAL-^R^ECO1* (SN41) in galactose-free medium, and *PDS5-AID* (SN80) treated with IAA. Cells were arrested at metaphase by benomyl treatment. See Figure S1 for details on culture conditions of *GAL-^R^ECO1* and *PDS5-AID*. The y-axis indicates normalized fold enrichment (nFE), or ChIP/input ratio normalized by spike-in control, with the range in brackets. CEN, centromere. The identified peaks are indicated by red horizontal bars, and the number of the peaks in the genome is shown in parentheses. **(B)** Heatmap of Scc1 and Scc2 ChIP-seq nFE in the indicated strains. 10kb-surrounding regions of the cohesin binding sites in wt (excluding ones less than 5 kb to the centromeres) are depicted. Regions are sorted in descending order of Scc2 nFE in Δ*wpl1* Δ*eco1*. **(C)** (Top) The scatter plot showing Scc2 nFE and the distance from the centromeres at each non-centromeric cohesin binding site in wt. Overlay representation of all chromosomes. The cohesin sites are divided into four groups according to their Scc2 FE and depicted by different colors. (Bottom) The kernel density estimation plot for the cohesin sites in each group. **(D)** Aggregated ChIP-seq profiles of Scc1 and Scc2 in wt and Δ*wpl1* Δ*eco1* for each group of the cohesin sites. The profiles are centered at the summit of the Scc1 peaks. Bold line, mean; shaded area, 95% confidence interval. See also Figure S1-S5.

The co-localization of Scc2 with Scc1 required simultaneous deletion of *WPL1* and *ECO1* (Figure 1A). It was not observed in Δ*wpl1* or the cells where Eco1 was depleted by promoter shut-off (*GAL-^R^ECO1*) (Yoshimura et al., 2021) (Figure 1A and S4). In the Eco1 depletion experiment, cells were allowed to progress through S phase in the absence of Eco1 and arrested in metaphase (Figure S1C). Scc2 co-localization was not seen also in the metaphase cells where Pds5 was acutely degraded by the auxin-inducible degron (AID) system (*PDS5-AID*) (Figure 1A and S1B). We compared the genome-wide binding profile of Scc1 and Scc2. Heatmap showing the nFE value around all cohesin binding sites on chromosome arms revealed that most of the cohesin sites were co-occupied with little, if any, Scc2 in wt, Δ*wpl1*, *GAL-^R^ECO1*, and *PDS5-AID* (Figure 1B). In contrast, Scc2 was found to accumulate at a majority of the same cohesin sites in Δ*wpl1 Δeco1*, although the amount of Scc2 binding was not sufficient to be called as a peak by the peak-calling algorithm for most of the sites (Figure 1B). The binding location of cohesin (Scc1) itself showed little difference between wt and Δ*wpl1* Δ*eco1*, though the binding intensity was reduced in general in Δ*wpl1* Δ*eco1*. ChIP-qPCR also confirmed the co-localization of Scc2 at the cohesin sites exclusively in Δ*wpl1* Δ*eco1* (Figure S3B).

We noticed that the intensity of Scc2 binding at the cohesin sites in Δ*wpl1* Δ*eco1* varied from site to site. To elucidate the nature of the strong Scc2 binding sites, the cohesin sites were equally divided into four classes based on Scc2 ChIP-seq nFE at each site (Q1 to Q4 in order of increasing nFE). The cohesin sites with the highest Scc2 binding (class Q4) were found to be enriched in regions less than 100 kb from the centromeres (Figure 1C). As mentioned above, Δ*wpl1* Δ*eco1* mutation caused a reduction of Scc1 ChIP-seq peak height. The ChIP-seq profiles averaged over the cohesin sites in each class indicated that the degree of the reduction in Scc1 binding was more severe in classes Q1 and Q2, where Scc2 co-localization was less prominent (Figure 1D).

Pds5 is a mutually exclusive antagonist of Scc2 in association with cohesin (Kikuchi & al., 2016; Petela et al., 2018). We investigated the chromosomal binding of Pds5 in Δ*wpl1* Δ*eco1* strain by spike-in ChIP-seq and ChIP-qPCR. The results consistently showed that the binding of Pds5 at the cohesin sites was significantly decreased in Δ*wpl1* Δ*eco1* strain (Figure S5), further verifying the increased Scc2 binding at the cohesin sites.

### Scc2 co-localization with cohesin in Δ*wpl1* Δ*eco1* is specific to metaphase

We conducted Scc1 and Scc2 ChIP-seq in S phase cells, where the establishment of sister chromatid cohesin was in progress. The cells were arrested in G1 phase by α-factor, followed by release into an α-factor-free medium for 30 min, to synchronize them in S phase (Figure 2A). Scc1 ChIP-seq profiles were similar between S and metaphase, though the binding was less intense, and the number of the detected peaks was decreased in S phase (Figure 2B, C). In wt cells, Scc2 binding profiles were similar between S and metaphase, with the only difference being that the binding at the centromere regions was less prominent in S phase (Figure 2C). Contrarily, in Δ*wpl1* Δ*eco1* cells, the co-localization of Scc2 at the cohesin sites was observed only in metaphase; the profile in S phase was almost identical to that in wt, and no increase in Scc2 binding at the cohesin sites was observed (Figure 2C, D). Hence, the co-localization of Scc1 and Scc2 in Δ*wpl1* Δ*eco1* was specific to metaphase. This result implies that the co-localization of Scc1 and Scc2 in Δ*wpl1* Δ*eco1* is related to the function of cohesin outside sister chromatid cohesion.

**Figure 2.**
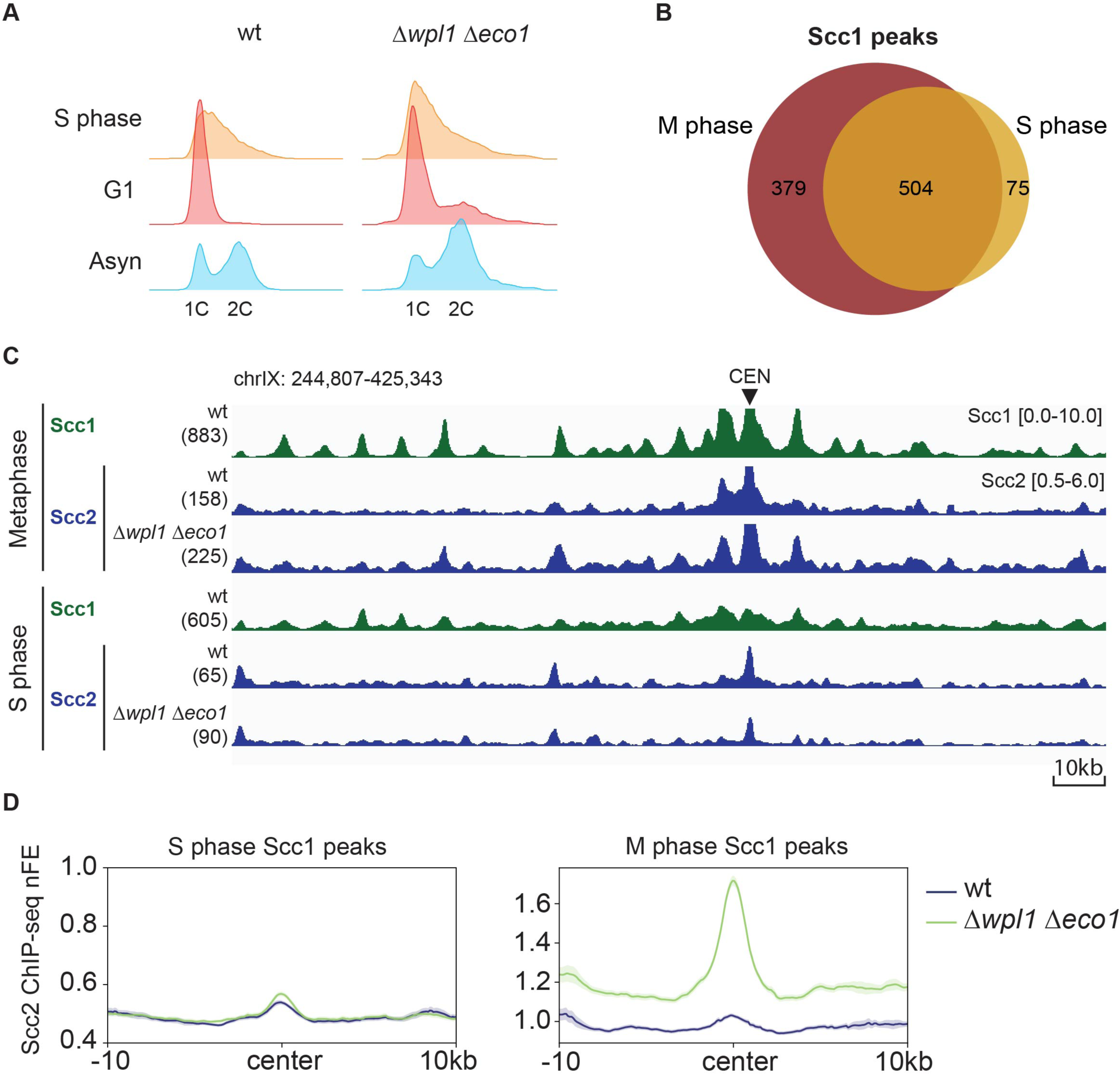
Scc2 colocalization with cohesin in Δ*wpl1 Δeco1* is specific to metaphase. **(A)** Cell cycle monitoring by flow cytometry. Asyn, asynchronous. Cells were synchronized in S phase by releasing G1-arrested cells for 30 mins in α-factor-free medium. The used strains were the same as Figure 1. **(B)** Venn diagram illustrating the overlap of Scc1 peaks in S phase and M phase. **(C)** Calibrated ChIP-seq profiles of Scc1 and Scc2 in the indicated conditions. **(D)** Aggregated ChIP-seq profiles of Scc2 in S-and M-phase wt cells. 10-kb surrounding regions of the non-centromeric cohesin sites in wt were averaged and depicted. Bold line, mean; shaded area, 95% confidence interval. Note that the y-axis ranges are different between the two plots.

### Δ*wpl1* Δ*eco1* promotes the extension of pericentromeric DNA loops

Since Scc2 binding confers the loop extrusion activity to cohesin, we investigated genome DNA folding of Δ*wpl1* Δ*eco1* mutant cells using Micro-C technique. Micro-C, an improved version of Hi-C, can reveal chromosome folding with higher resolution (Hsieh et al., 2015) and indeed enabled us to detect each loop clearly without the need of ‘pile-up’ analysis in yeast chromosomes. In wt, consistent with previous study (Costantino et al., 2020), short DNA loops of several tens of kb, which connect adjacent cohesin binding sites, were observed along the entire length of the chromosomes (Figure 3A, B). These short DNA loops largely disappeared in Δ*wpl1* Δ*eco1* mutant. Similar disappearance of DNA loops along chromosome arms was described in Pds5*-*depleted cells previously (Bastié et al., 2022; Costantino et al., 2020). In Δ*wpl1* Δ*eco1*, we observed novel, longer DNA loops that connect a centromeric region and particular sites on the same chromosome (Figure 3A, B). The loops detected in Δ*wpl1* Δ*eco1* were smaller in number (159 compared with 240 in wt), but longer in length (mean loop length of 67 kb in Δ*wpl1* Δ*eco1* compared with 20 kb in wt) (Figure 3C). Consistently, the contact-versus-distance decaying curve and its derivative (Figure 3D) confirmed the decrease in short-range contacts (< 50kb) and increase in long-range contacts (> 50kb). Besides loop length, the loop interaction was also more intense in Δ*wpl1* Δ*eco1* mutant than wt, which was revealed by the aggregate peak analysis (APA) plots over the cohesin-positive loops (Figure 3E).

**Figure 3.**
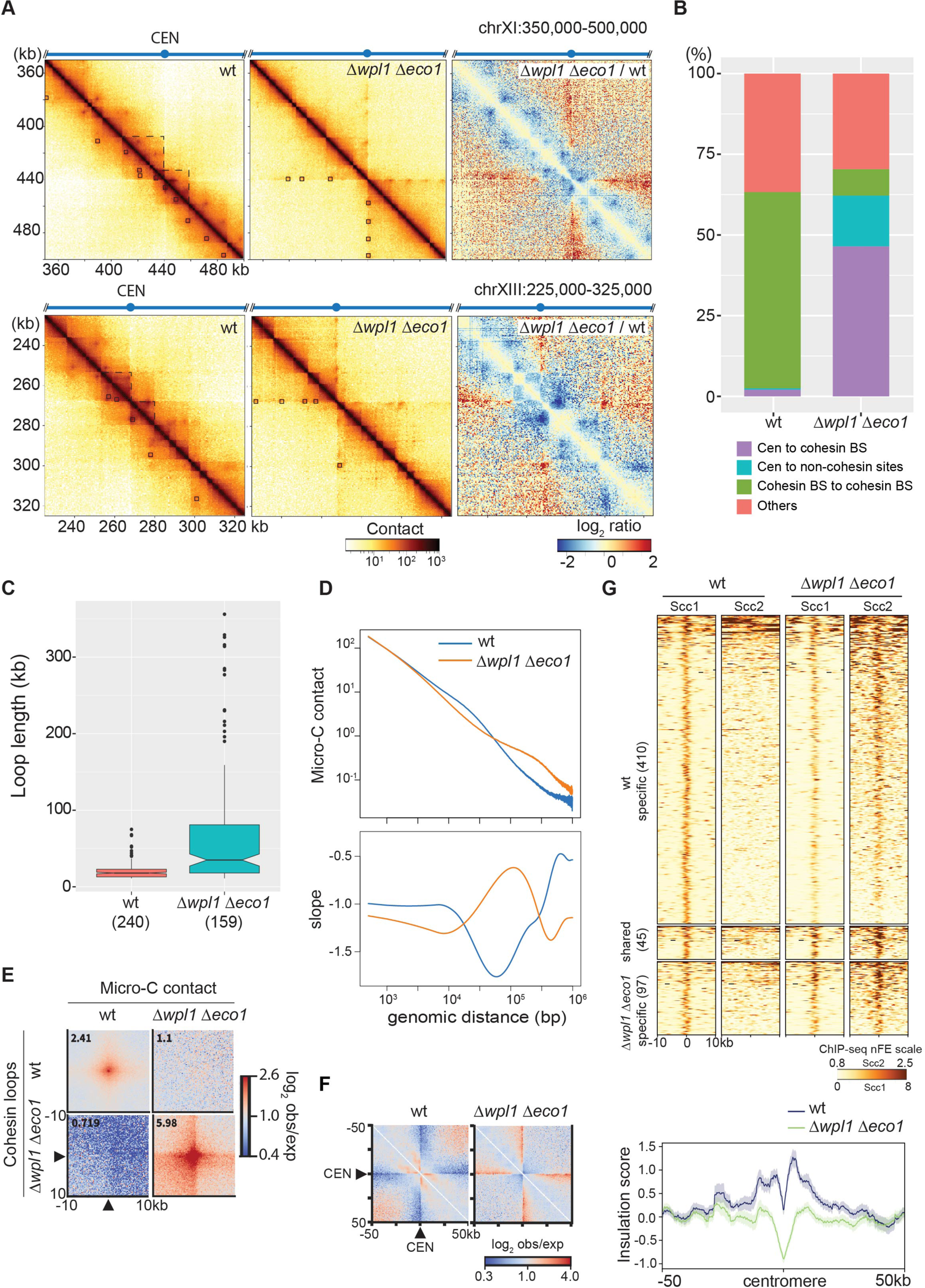
Δ*wpl1* Δ*eco1* promotes extension of pericentromeric DNA loops. **(A)** Micro-C contact maps in wt (SKY001), Δ*wpl1* Δ*eco1* (KT110), and log2-ratio between Δ*wpl1* Δ*eco1* and wt. Cells were arrested at metaphase by benomyl treatment. Bin size is 500 bp. Squares in the contact maps indicated loops called by HICCUPs. Dashed lines mark the chromosomal domains adjacent to centromeres. Two representative regions in chromosomes XI and XIII are shown. The number of valid pairs in each sample was normalized to 115 million. CEN, centromere. **(B)** Proportion of the detected loops in each indicated category. Cen, centromere; cohesin BS, cohesin binding site. **(C)** Length distribution of cohesin loops (loops connecting between cohesin BSs) in wt and Δ*wpl1* Δ*eco1*. **(D)** Contact-versus-distance decaying curves of the Micro-C contact matrix in wt and Δ*wpl1* Δ*eco1,* and their first derivatives (slope). **(E)** Average contact frequency between cohesin loop anchors in wt and Δ*wpl1* Δ*eco1*. Observed/expected ratio of the contact frequency around the off-diagonal peaks connecting two cohesin loop anchors found in wt or Δ*wpl1* Δ*eco1* were averaged and plotted. The number in the top-left corner of each plot indicates the average enrichment score of the 3 × 3 central pixels. **(F)** (Left) Average contact frequency around the centromeres in wt and Δ*wpl1* Δ*eco1*. Black triangles indicate the centromere position. (Right) Averaged insulation score around the centromeres in wt and Δ*wpl1* Δ*eco1*. Bold line, mean; shaded area, 95% confidence interval. **(G)** Heatmaps of Scc1 and Scc2 ChIP-seq nFE in wt and Δ*wpl1* Δ*eco1*. 10-kb surrounding regions of the cohesin-bound loop anchors specific to wt, specific to Δ*wpl1* Δ*eco1*, and shared between wt and Δ*wpl1* Δ*eco1* are depicted. The number of sites for each group is shown in parentheses.

Δ*wpl1* Δ*eco1* also caused changes in the chromatin compartmentalization around centromeres in Δ*wpl1* Δ*eco1*, as shown in Figure 3A. First, self-interactions within a region that is about 10-20 kb in size and adjacent to the centromere, which is prominent in wt, became weaker in Δ*wpl1* Δ*eco1*. Second, the centromere-adjacent regions on both chromosome arms became more insulated in Δ*wpl1* Δ*eco1*. These points were verified by the normalized APA plot of contact frequency on all centromeres (Figure 3F, left). The insulation score plot around the centromeres (Figure 3F, right) also confirmed the disappearance of inter-arm interaction in Δ*wpl1* Δ*eco1*; a low insulation score at the centromere indicates less-than-expected interaction across it. Together, the data indicates that Δ*wpl1* Δ*eco1* caused drastic changes in higher-order chromosome folding, including the expansion of pericentromeric loops.

### Novel loop anchors in Δ*wpl1* Δ*eco1* are associated with cohesin and Scc2 binding

As shown in Figure 3B, most of the DNA loops were anchored at cohesin binding sites. The number of the loop anchor sites with cohesin binding (excluding the centromere regions) was 455 and 142 in wt and Δ*wpl1* Δ*eco1*, respectively, and 45 of them were shared. We examined the binding of Scc2 at these sites by plotting the heatmaps of Scc2 ChIP-seq profile in a 10-kb flanking region centered at the cohesin-bound loop anchors (Figure 3G). In wt, Scc2 was hardly found at cohesin-bound loop anchors. Contrarily, in Δ*wpl1* Δ*eco1* cells, Scc2 binding was observed at the anchor sites. The binding was more intense in general at the anchors detected in Δ*wpl1* Δ*eco1*. These data suggest that the co-localization of cohesin and Scc2 is coupled with the emergence of novel DNA loops in Δ*wpl1* Δ*eco1*.

### DNA loop extension is blocked by centromere-oriented long genes

The centromere-originated loops in Δ*wpl1* Δ*eco1* reached the regions more than 100 kb from the centromere. This is in sharp contrast to wt cells, where loops reach only a few tens of kb from the centromeres. As mentioned above, these loops typically ended at the cohesin binding sites, but only a fraction of the cohesin binding sites served as the anchors. To clarify the nature of the anchor sites, we divided the cohesin sites in the genome into two groups: anchor (142) and non-anchor (813) cohesin sites, which exhibited or did not exhibit the interaction with the centromere on the same chromosome, respectively (Figure 4A). We found that the binding of both cohesin and Scc2 was more intense at the anchor cohesin sites, compared with the non-anchor cohesin sites (Figure 4B). Consistently, when the cohesin binding sites were grouped based on the level of co-localized Scc2 as in Figure 1D, the contact frequency between the cohesin site and the centromere was getting more intense for the cohesin sites with more Scc2 binding (Figure S6). We also noticed that Δ*wpl1* Δ*eco1* resulted in a substantial increase in cohesin binding of cohesin at the centromeres (Figure 4B).

**Figure 4.**
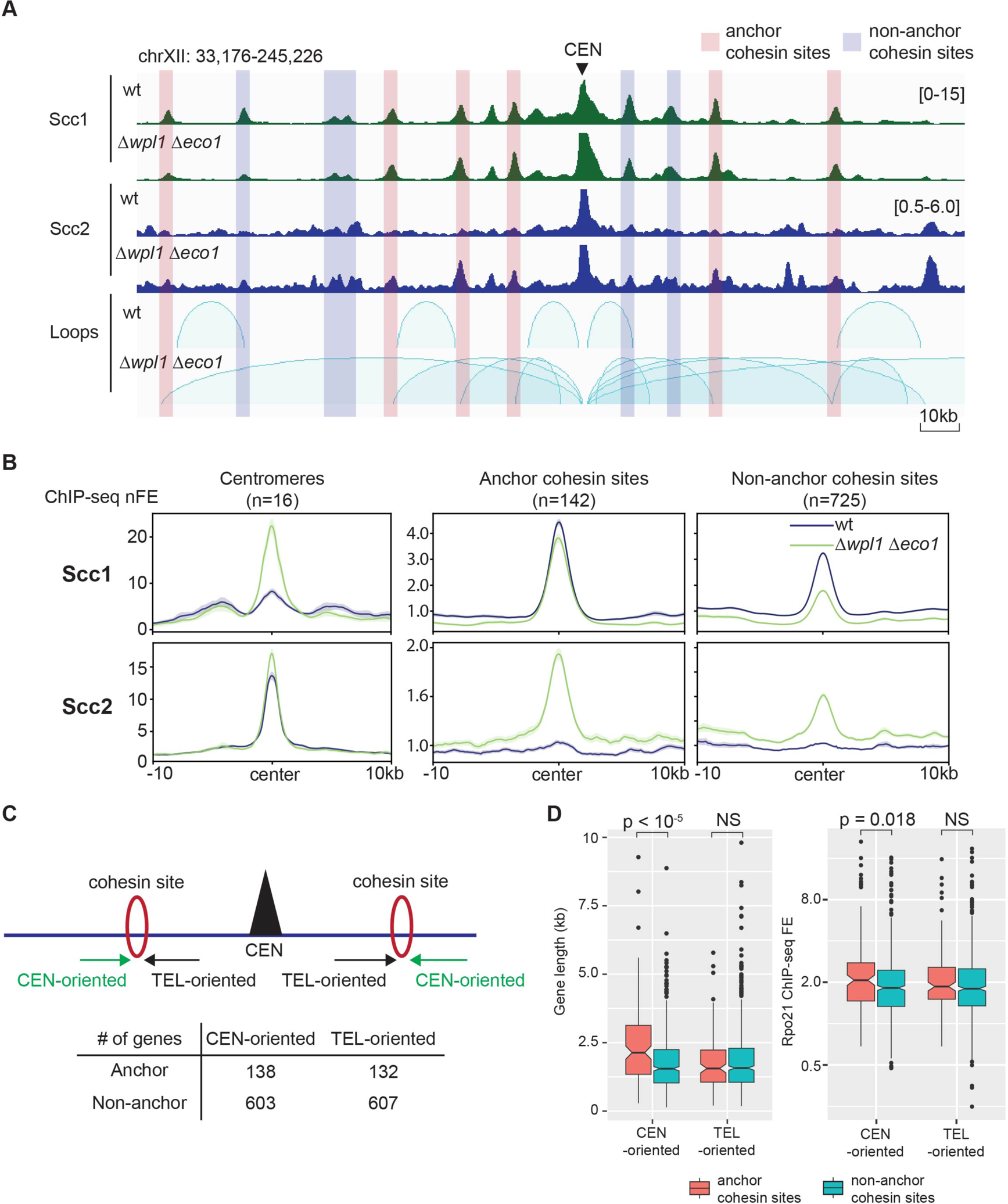
DNA loop expansion in Δ*wpl1* Δ*eco1* is impeded by long centromere-oriented genes. **(A)** ChIP-seq profile of Scc1 and Scc2 alongside loops called by HICCUP in wt and Δ*wpl1* Δ*eco1* arrested at metaphase. The cohesin sites are categorized into anchor or non-anchor sites depending on their overlapping with anchors of centromere-originated loops in Δ*wpl1* Δ*eco1*. **(B)** Aggregated plots of Scc1 and Scc2 ChIP-seq nFE in wt and Δ*wpl1* Δ*eco1*. 10-kb surrounding regions around the centromeres, anchor cohesin sites, and non-anchor cohesin sites are depicted. Bold line, mean; shaded area, 95% confidence interval. **(C)** Labeling of the genes adjacent to the cohesin binding sites. CEN-oriented genes, genes distal to the centromere (CEN) and transcribed toward it. TEL-oriented genes, genes proximal to CEN and transcribed away from it. Note that most of the cohesin sites are located in convergent intergenic regions. The number of genes in each category is shown in the table. **(D)** Boxplot comparison of gene length and Rpo21 ChIP-seq FE between CEN-and TEL-oriented genes. The numbers above the plots are p-values (Mann-Whitney *U* test, two-tailed). NS, non-significant difference (p > 0.05). See also Figure S6 and S7.

Gene transcription can be an obstacle to loop expansion (Banigan et al., 2023; Jeppsson et al., 2022; Paldi et al., 2020). We, thus, addressed whether the neighboring genes of the anchor cohesin sites display any specific characteristics. As reported (Lengronne et al., 2004), most of the cohesin binding sites in yeast were located in the intergenic regions (IGRs) between two convergently oriented genes. We observed that genes located on the side distal to the centromere and transcribed towards the centromere (referred to as CEN-oriented genes, as illustrated in Figure 4C) were longer when associated with anchor cohesin sites compared to non-anchor cohesin sites (Figure 4D, left). Additionally, the transcriptional activity of these CEN-oriented genes, measured by Rpo21 ChIP-seq FE, was significantly higher at anchor cohesin sites (Figure 4D, right). In contrast, genes situated proximal to the centromere and transcribed towards the telomeres (referred to as TEL-oriented genes, Figure 4C) did not exhibit any differences in either length or transcriptional activity between anchor and non-anchor cohesin sites (Figure 4D). These data strongly indicate that length and transcription activity of the CEN-oriented genes plays a substantial role in the determination of the loop location in *Δwpl1 Δeco1*.

Besides loops extruded from centromeres, we also occasionally observed the “stripes” projecting from several loci on chromosome arms in Δ*wpl1* Δ*eco1* (Figure S7, left). Examining those loci revealed that the neighboring genes are either exceptionally long (ex. *GCN1*, *MYO1*, and *FAS1*) or strongly transcribed (ex. *PDR5*) (Figure S7, right). The loop anchors were coincident with the 3’ end of these genes and associated with high-level cohesin binding. The loops always extend only toward the downstream direction of the genes. We speculate that these genes act as strong, direction-dependent barriers to the expansion of the loops initiated somewhere downstream of the gene, thereby leading to the stripe pattern in the Micro-C contact map.

### Synergistic effect of Δ*wpl1* and *Δeco1* in loop extension

Previous studies have reported that the pericentromeric loops are also extended in Δ*wpl1* single deletion mutant and cells where Pds5 is acutely depleted by auxin-inducible degron (*PDS5-AID*) (Bastié et al., 2022; Costantino et al., 2020; Dauban et al., 2020). We conducted Micro-C analysis in Δ*wpl1* and Pds5-depleted cells in parallel with Δ*wpl1* Δ*eco1* to compare the characteristics of DNA loops. All the strains were arrested in metaphase. Vehicle-treated *PDS5-AID* strain, as a control, exhibited only short DNA loops along the chromosomes, like wt cells (Figure 5A, B). Δ*wpl1* and IAA-treated *PDS5-AID* cells demonstrated the extension of the centromere-originated loops as reported previously but to a lesser extent than Δ*wpl1* Δ*eco1* double deletion mutant (Figure 5A, B). The number and length of the centromere-originated loops were also moderately increased in Δ*wpl1* and IAA-treated *PDS5-AID* but more prominently in Δ*wpl1* Δ*eco1* (Figure 5D, left and middle). To quantify the extension of centromere-originated loops genome-wide, we grouped the cohesin binding sites based on the distance from the centromere and depicted an aggregated plot for the interaction between the cohesin sites in each group and the corresponding centromere. The interaction became weaker as the cohesin sites were further away from the centromere (Figure 5B). The extent of the interaction in Δ*wpl1* and Pds5-depleted cells reached farther than in wt cells but remained much shorter than in Δ*wpl1* Δ*eco1* double deletion mutant (Figure 5B). The contact-versus-distance decaying curve and its derivative also confirmed an increase in long-range contacts of Δ*wpl1* and IAA-treated *PDS5-AID* but not as much as of Δ*wpl1* Δ*eco1* (Figure 5C). Different from Δ*wpl1* Δ*eco1*, Scc2 co-localization was not detected at the anchor cohesin sites in Δ*wpl1* or IAA-treated *PDS5-AID* cells (Figure 5D, right).

**Figure 5.**
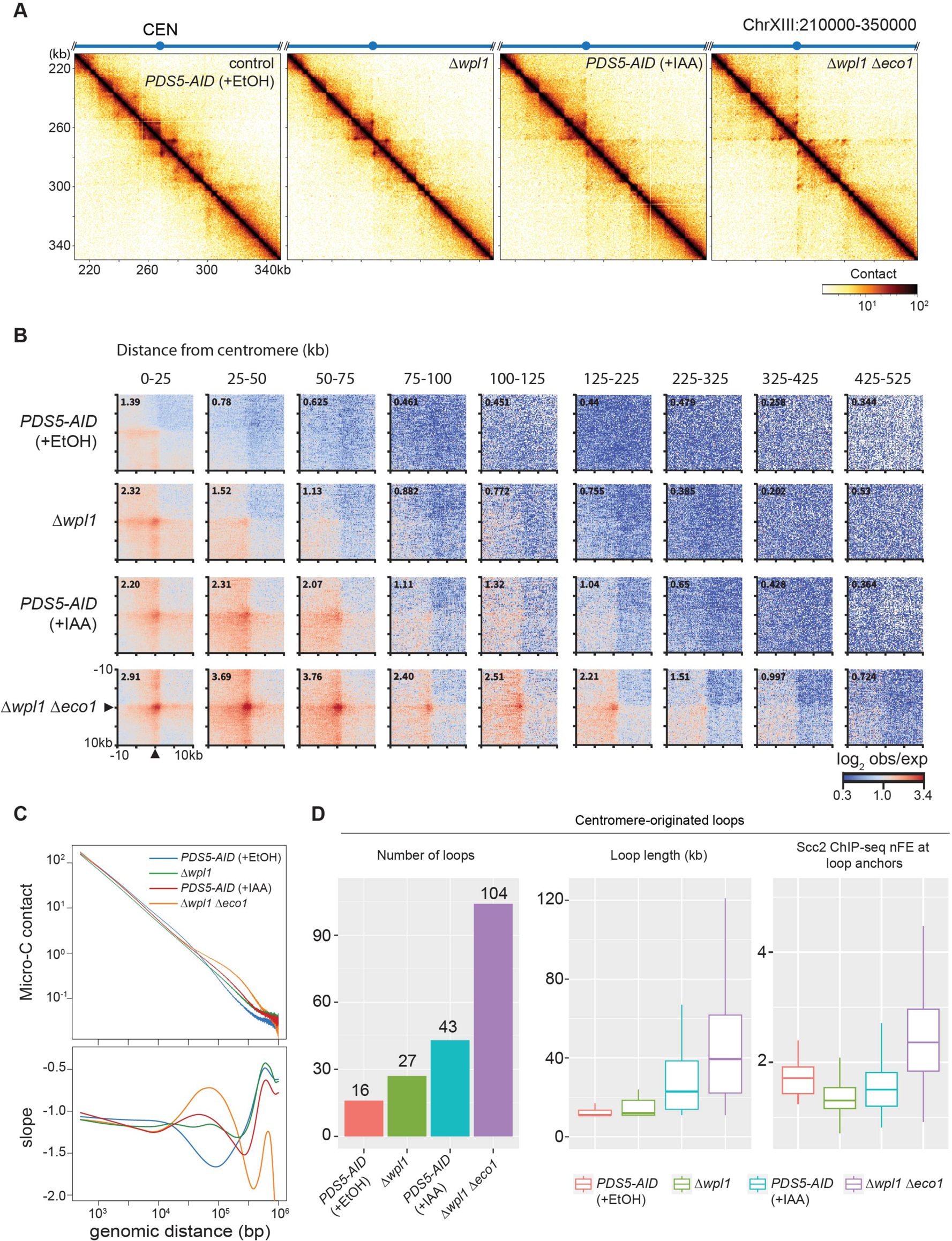
Synergetic effect of Wpl1 and Eco1 depletion on loop extension. **(A)** Micro-C contact maps in *PDS5-AID* (SN80) treated with vehicle (+EtOH) or IAA, Δ*wpl1* (KT127), and Δ*wpl1* Δ*eco1* (SN54) cells arrested at metaphase. Vehicle-treated *PDS5-AID* serves as a control. *PDS5-AID* cells were cultures as in Figure S1B. Contact maps were computed on 500 bp-resolution data. The number of valid pairs in each sample was normalized to 42 million. **(B)** Average contact frequency (represented as observed/expected ratio) between a centromere and a cohesin site on the same chromosome in the indicated strains arrested at metaphase. The centromere-cohesin site pairs were grouped by the distance between the two sites. **(C)** Contact-versus-distance decaying curves of the contact matrix in the indicated strains, and their first derivatives (slope). **(D)** Comparison of the centromere-originated loops identified in the indicated samples. (Left) The number of the loops. (Middle) Length distribution. (Right) Scc2 ChIP-seq nFE at the non-centromeric anchor sites. Outliers were not depicted in the box plots.

### Centromere-originated loops in Δ*wpl1* Δ*eco1* are dependent on Scc2

The co-localization of Scc2 and cohesin in Δ*wpl1* Δ*eco1* implied that the extended pericentromeric DNA loops were generated by the Scc2-dependent loop extrusion activity of cohesin. To verify this, we investigated chromosome folding in the cells where Scc2 was acutely depleted by the AID system. First, we verified the efficacy of AID-induced Scc2 depletion. In budding yeast, cohesin is almost absent in α-factor-arrested G1 cells and loaded onto chromosomes during late G1–S phase. When G1-arrested cells were released and re-arrested at metaphase in the presence of IAA, we observed very little cohesin loading at all the cohesin sites in the genome (Figure S8A, +IAA(S)). This clearly indicates that the AID system successfully depleted Scc2, which is required for cohesin loading onto chromosomes, to very low level. We, then, tested another condition, where Scc2 was depleted after metaphase (Figure 6A and S8A, +IAA). In this condition, cohesin loading onto the chromosome arms was only marginally attenuated, indicating that the latter experimental scheme allows the suppression of Scc2 activity in metaphase without affecting sister chromatid cohesion formation in S phase. Scc1 binding level at the centromeres was greatly reduced in the second condition, particularly in Δ*wpl1* Δ*eco1* (Figure S8A, right). This is consistent with the fact that centromeres are active cohesin loading sites in metaphase cells without spindle tension (Ocampo-Hafalla et al., 2007).

**Figure 6.**
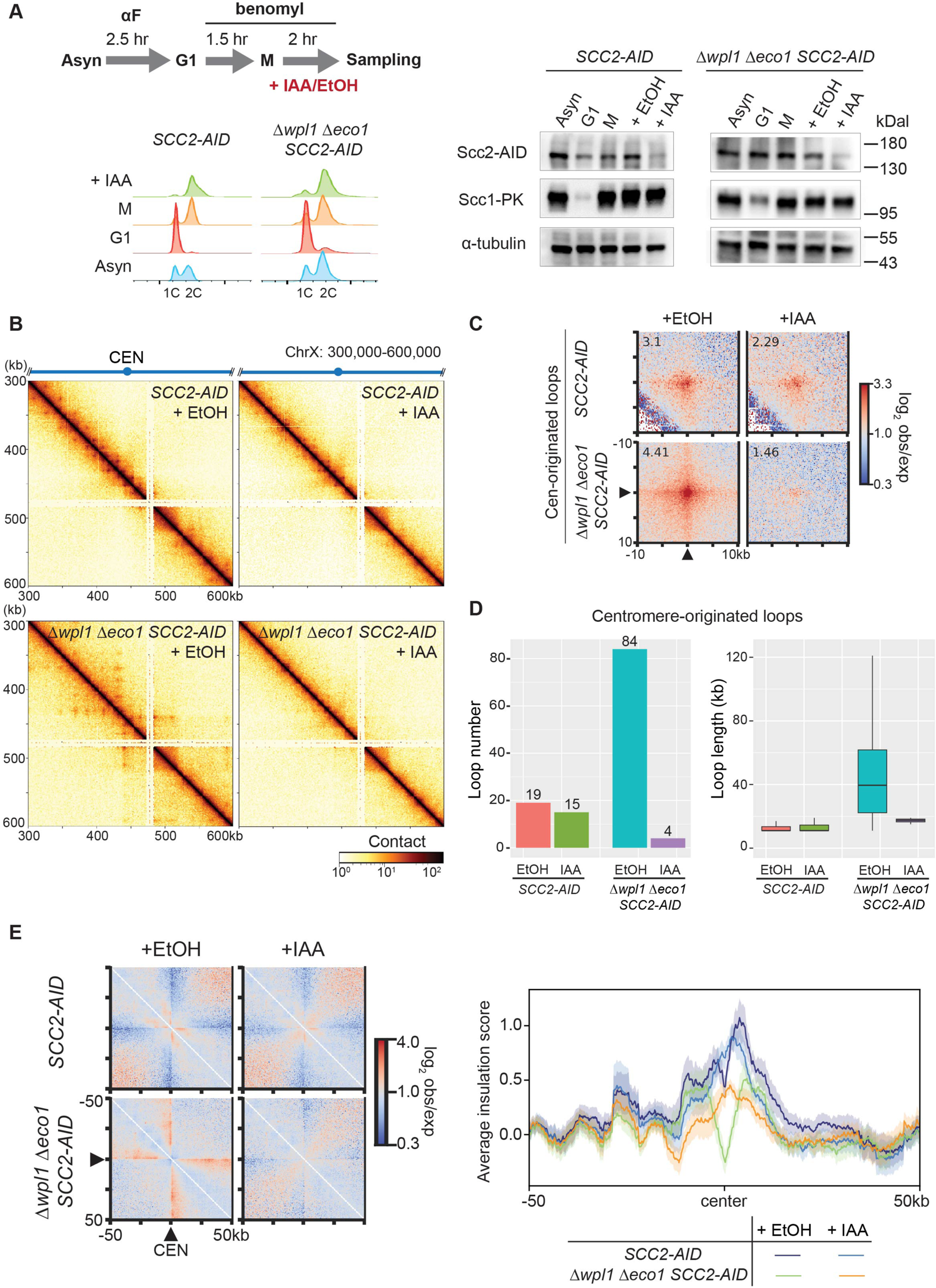
**Acute depletion of Scc2 eliminates cohesin-mediated extended loops in Δ*wpl1* Δ*eco1*** **(A)** (Left) Schematic representation of the experimental protocol used to arrest cells in metaphase and induce rapid degradation of Scc2-AID. (Right) Western blot to verify Scc2-AID depletion. **(B)** Micro-C contact maps in *SCC2-AID* (SN75) and Δ*wpl1* Δ*eco1 SCC2-AID* (SN74) strains treated with vehicle (EtOH) or IAA at the resolution of 1 kb. The number of valid reads in each sample was normalized to 28 million. **(C)** Average contact frequency for the centromere-originated loops detected in vehicle-treated *SCC2-AID* and Δ*wpl1* Δ*eco1 SCC2-AID* strains (+EtOH). The averaged contact frequency for the same locus-pairs in IAA-treated condition (+IAA) was shown side by side for comparison. **(D)** The number and length of centromere-originated loops detected in the indicated conditions. Outliers were omitted from the box plot. **(E)** (Left) Average contact frequency around the centromeres. Black triangles indicate the centromere position. (Right) Averaged insulation score around the centromeres in the indicated condition. Bold line, mean; shaded area, 95% confidence interval. See also Figure S8 and S9.

We conducted Micro-C analysis of the cells where Scc2 was depleted after metaphase arrest. As a control, we analyzed Δ*wpl1* Δ*eco1 SCC2-AID* strain (strain SN74) in the absence of IAA, which revealed some changes in the contact map compared with the untagged Δ*wpl1* Δ*eco1* strain (KT110) (Figure S9A). Of particular, loops connecting the cohesin sites on chromosome arms were detected in addition to the centromere-originated extended loops, and the number of detected loops was larger in SN74 (Figure S9B, left). These differences were reproducibly observed, implying that the difference was not due to experimental variability. Nevertheless, the overall trend in chromosome folding change was identical between the two strains; both KT110 and SN74 exhibited significantly longer loops than wt (Figure S9B, right) and a similar decrease in short-range contacts accompanied by an increase in long-range contacts (Figure S9C). In addition, average contact maps around centromeres in SN74 portrayed the loss of centromere-neighboring domains and the emergence of long-range centromere-originated interactions, like KT110 (Figure S9D). We hence concluded that untagged Δ*wpl1* Δ*eco1* and Δ*wpl1* Δ*eco1 SCC2-AID* strains shared the same characteristic in centromere-involved chromosome folding and used SN74 strain for subsequent experiments.

The Micro-C contact map revealed that depletion of Scc2 in metaphase-arrested Δ*wpl1* Δ*eco1* cells resulted in the disappearance of the centromere-originated loops (Figure 6B); they were greatly reduced in both contact frequency, number, and length (Figure 6C, D). The same Scc2 depletion showed only a modest impact on the centromere-originated loops in wt (Figure 6C). Scc2 depletion also affected the additional arm-to-arm loops found in SN74 strain, because off-diagonal APA plots for all cohesin-bound loop anchors showed that the average genome-wide contact frequency was drastically reduced in Δ*wpl1* Δ*eco1* (from 3.14 to 1.61) (Figure S9B). On-diagonal APA analysis and average insulation score profile around centromeres indicate that Scc2 depletion in metaphase resulted in the loss of the insulating feature of centromeres in Δ*wpl1* Δ*eco1* (Figure 6E). Taken together, we conclude that the Scc2 activity in metaphase is required for the formation of cohesin-mediated loops and the insulation across the centromeres seen in Δ*wpl1* Δ*eco1*.

### Extended pericentromeric loops impede chromosome segregation process

We explored how the extension of the pericentromeric DNA loops affects the function of the kinetochore. Wt and Δ*wpl1* Δ*eco1* strains possessing *SCC2-AID* were arrested in metaphase with benomyl and released from the arrest, followed by DAPI-based monitoring of nuclear division (the detailed experimental scheme is in Figure 7A, left). In wt, anaphase or telophase cells, or cells with segregated nuclei, started to appear 40 mins after the release (Figure 7A, right). In contrast, Δ*wpl1* Δ*eco1* cells exhibited a delay in the nuclear division because the proportion of the anaphase or telophase cells was significantly smaller than wt at 40 min after the release. Consistently, the proportion of large-budded cells with a single nucleus remained high, compared with that in wt, throughout the course of the experiment. Of note, the delay in nuclear division was almost completely rescued by Scc2 depletion in metaphase. Scc2 depletion in wt had little impact on the timing of nuclear division. This result suggests that the extended DNA loops around centromeres, of which the formation is dependent on Scc2, impede the smooth progression of nuclear division.

**Figure 7.**
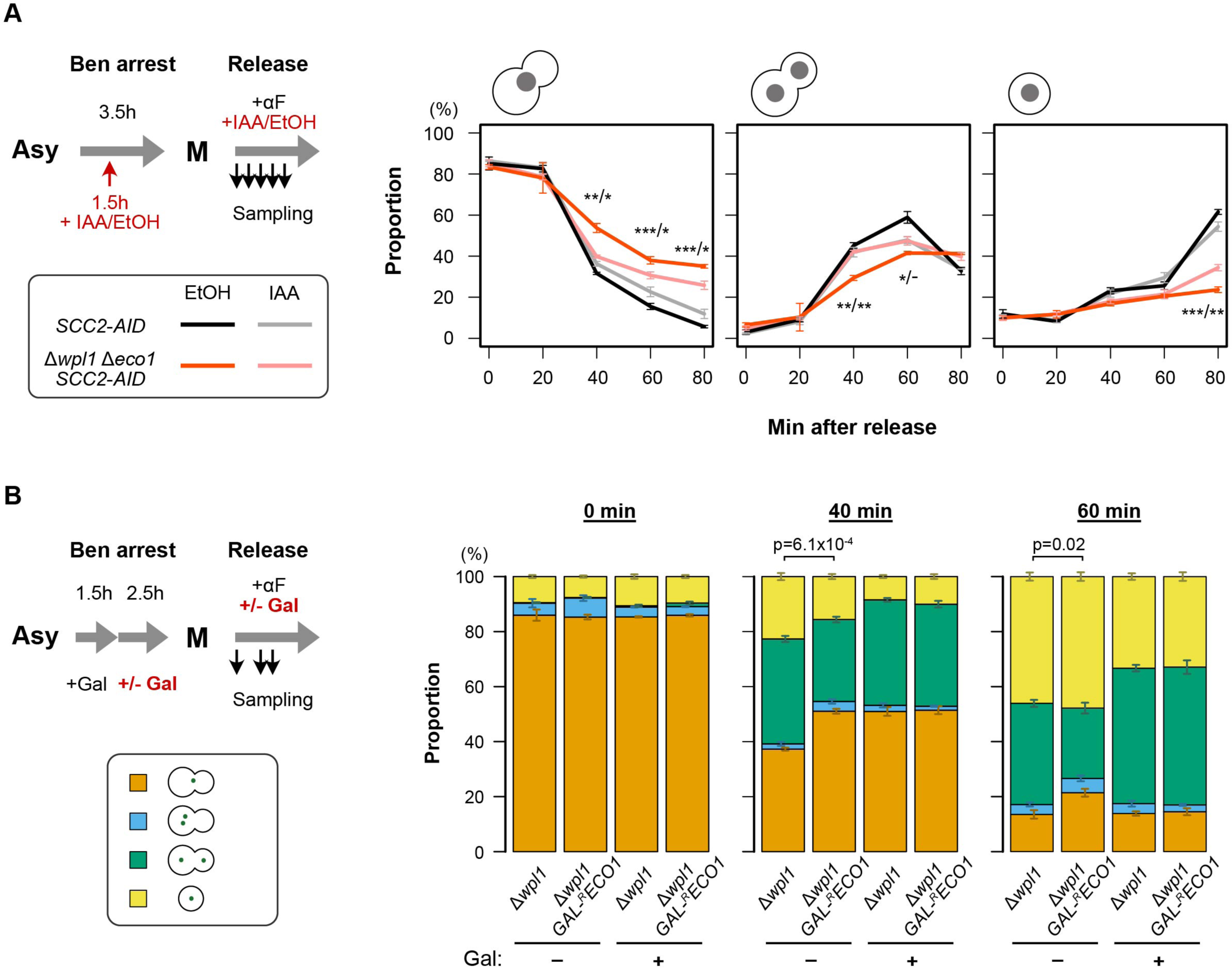
Expanded pericentromeric loops impeded the smooth progression of chromosome segregation. **(A)** (Left) Schematic representation of the experimental protocol to make cells proceed synchronously through mitosis in the absence or presence of Scc2. *SCC2-AID* (SN75) and Δ*wpl1* Δ*eco1 SCC2-AID* (SN74) were analyzed. Ben, benomyl; αF, α-factor. (Right) Proportion of budded cells with one nucleus, budded cells with two nuclei, and unbudded cells with one nucleus (from left to right). The x-axis shows time points (in minutes) after release from benomyl arrest. The mean of three biological replicates, in each of which > 150 cells were counted, are shown. Error bar, SEM. Two-tailed t-test was conducted to evaluate the statistical significance of the difference (i) between Δ*wpl1* Δ*eco1 SCC2-AID* and *SCC2-AID* in EtOH-treated condition, and (ii) between EtOH-and IAA-treated conditions of Δ*wpl1* Δ*eco1 SCC2-AID*. The two test results were displayed with a slash symbol in between. ***, p < 0.0001; **, p < 0.01; *, p < 0.05;-, p < 0.1. **(B)** (Upper left) Schematic representation of the experimental protocol to make cells proceed synchronously through mitosis in the absence or presence of Eco1. Δ*wpl1* (ST258) and Δ*wpl1 GAL-^R^ECO1* (ST722) were analyzed. Gal, galactose. (Lower left) Categorization of cells based on morphology and *URA3*-GFP foci. (Right) The proportion of cells in each category. Cells were collected at 0, 40 or 60 minutes after the release from benomyl-arrest. The means of three biological replicates, in each of which > 150 cells were counted, are shown. Error bar, SEM. Two-tailed t-test was conducted to evaluate the statistical significance of the difference in the fraction of budded cells with a single GFP dot (orange) between Δ*wpl1* and Δ*wpl1 GAL-^R^ECO1* in the absence of galactose, and the calculated p-values were shown.

We conducted another experiment to test this notion. Δ*wpl1 GAL-^R^ECO1* strain was arrested in metaphase with benomyl, and the medium was changed to galactose-free YPD to repress Eco1 expression. Then, the cells were released from the arrest, and chromosome segregation was monitored by the TetO/TetR-GFP mark at *URA3* locus (experimental scheme in Figure 7B, left). At metaphase, Δ*wpl1* Δ*eco1* shows a higher proportion of cohesion-defective cells than Δ*wpl1* (Sutani et al., 2009). We, however, observed little difference in the proportion of cohesion-defective cells between Δ*wpl1 GAL-^R^ECO1* and the control Δ*wpl1* at the timing of the release (0 min). This indicates that the cells passed through S phase with functional Eco1, and sister chromatid cohesion was established with little defect. At 40 and 60 mins after the release, Δ*wpl1 GAL-^R^ECO1* showed a significantly higher proportion of the cells with unsegregated GFP focus than Δ*wpl1*, indicating a delay in mitotic progression (Figure 7B, right). Since the degree of the pericentromeric DNA loop extension was more prominent in Δ*wpl1* Δ*eco1* than Δ*wpl1*, this data also supports the notion that the extended pericentromeric loops are detrimental to timely chromosome segregation. As a control, we monitored cell division also in galactose-containing YPG medium. Δ*wpl1 GAL-^R^ECO1* and Δ*wpl1* stains showed no difference in mitotic progression. The progression was slower in YPG, which may be because galactose is a less preferred energy source.

## Discussion

In wild-type budding yeast, Scc2 shows no co-localization with cohesin on chromosome arms (Hu et al., 2011). We found that Scc2 co-localized with cohesin at cohesin binding sites in Δ*wpl1* Δ*eco1* strain, suggesting that Scc2 interacts with cohesin more stably in this condition. In the genome of the same strain, the binding of Pds5 to the cohesin sites was significantly attenuated. This is consistent with the notion that the interaction of Pds5 with cohesin is antagonistic to that of Scc2 and cohesin (Kikuchi & al., 2016; Petela et al., 2018). The co-localization of Scc2 and cohesin was not observed in Δ*wpl1* strain or in cells from which only Eco1 was depleted. Therefore, we conclude that Eco1 and Wpl1 function synergistically in the emergence of Scc2-associated cohesin on budding yeast chromosomes.

Structural and biochemical studies suggest that cohesin acetylated by Eco1 loses its affinity for Scc2 (Kaushik et al., 2023; Shi et al., 2020). This explains well the contribution of Eco1 to Scc2-cohesin co-localization in Δ*wpl1* Δ*eco1*. Wpl1 promotes dissociation of non-acetylated cohesin from chromosomes (Lopez-Serra et al., 2013). In fact, the height of the cohesin peak was reduced by 70% in Eco1-depleted cells compared with wt. This effect partially explains why Scc2 co-localized with cohesin was not observed in the Eco1-depleted cells. In addition, we speculate that Wpl1 may have a more direct role in the control of Scc2-cohesin interaction. Recent structural predictions by AlphaFold2 imply the possibility that Wpl1 interacts with both Pds5 and cohesin to form ternary complexes (Nasmyth et al., 2023). Δ*wpl1* single deletion mutant actually caused a substantial reduction of Pds5 binding at cohesin sites (Figure S5C). Wpl1 deletion is, thus, likely to weaken the Pds5-cohesin interaction, thereby promoting Scc2 binding to cohesin.

In mitotically arrested wild-type cells without the spindle, centromere-anchored DNA loops of ∼10 kb in length are observed (Paldi et al., 2020). These loops are thought to be formed by cohesin bound to the centromeres and catalyzing the DNA loop extrusion reaction. We observed centromere-anchored DNA loops of more than 100 kb in length in Δ*wpl1* Δ*eco1* strain. This is, to our knowledge, the first report of such a scale of loop size expansion in yeast. Most likely, Scc2-associated cohesin remains ATPase active and continuously exhibits the loop extrusion reaction, resulting in extended DNA loops. Strong genes transcribed toward the centromere could act as a permeable barrier to loop expansion, but cohesin that maintains activity is assumed to overcome the barriers one by one, allowing continuous loop expansion. This idea is supported by the observations that the extended pericentromeric loops were not detected after the removal of Scc2 in mitotic cells and that the amount of Scc2 co-localized with cohesin is inversely correlated with the distance from the centromere. Hence, we conclude that Eco1 and Wpl1 additively contribute to the size restriction of pericentromeric DNA loops.

In wild-type cells, re-orientation of the loop boundary genes impeded the establishment of chromosome bipolar attachment (Paldi et al., 2020). In this study, we demonstrated that Δ*wpl1* Δ*eco1* double deletion mutant exhibited a delay in mitotic chromosome segregation, revealing that *trans* factors also play a role in facilitating the smooth progression of chromosome segregation through the restriction of the pericentromeric DNA loop size. As suggested by Paldi *et al*., the expanded pericentromeric DNA loops may disrupt sister chromatid cohesion in wider chromosome regions, thereby disturbing chromosome segregation. In addition, the presence of the large DNA loops nearby the centromeres may hinder kinetochore-microtubule attachment directly. Δ*wpl1* Δ*eco1* mutant is highly sensitive to a microtubule depolymerization drug, benomyl (Sutani et al., 2009). The expanded pericentromeric DNA loops may be the mechanism behind this phenotype.

Eco1 is essential for the establishment of sister chromatid cohesion during S phase (Tóth et al., 1999). In this study, we observed that mitosis-specific depletion of Eco1 in Δ*wpl1* strain was sufficient to cause a delay in chromosome segregation similar to that seen in Δ*wpl1* Δ*eco1* strain. This implies that Eco1 must function in metaphase to restrict pericentromeric DNA loop expansion. Consistently, Eco1 reportedly remains active in cohesin acetylation even in metaphase-arrested cells (Rowland et al., 2009).

A very recent report revealed that purified yeast cohesin with Scc2-Scc4 exhibits DNA loop extrusion activity almost equivalent to that of human cohesin *in vitro* (Guérin et al., 2023). Our current study demonstrated that yeast cohesin is actually capable of forming large DNA loops of several hundred kb *in vivo*, comparable to those in human nuclei if Scc2-cohesin interaction is free from inhibitory regulation by Wpl1 and Eco1. In this context, it is notable that co-localization of Scc2 and cohesin was not observed in S phase Δ*wpl1* Δ*eco1* cells. Other factors may regulate cohesin specifically at unattached centromeres in metaphase cells so that it exhibits higher affinity to Scc2 and becomes competent to DNA loop extrusion. It will be interesting to explore further how the Scc2-cohesin interaction is regulated in a cell cycle-and context-dependent manner.

## Acknowledgments

We thank Dr. Kristian Jeppsson and all members of Shirahige laboratory for discussion. This work was supported by JST CREST grant number JPMJCR18S5, JSPS KAKENHI grant numbers JP20H05940, JP20H05933 and JP20H05686, and AMED ASPIRE-A grant number JP23jf0126003 (to K.S.); JSPS KAKENHI grant number JP21K06012 (to T.Su.).

## Author contributions

S.A.N., K.S., and T.Su. designed the study and interpreted the data. S.A.N. performed the experiments. S.A.N. and T.Sa. analyzed data. S.A.N. and T.Su. wrote the manuscript. S.A.N., T.Sa., K.S., and T.Su. reviewed and edited the manuscript.

## Methods

KEY RESOURCES TABLE.

KEY RESOURCES TABLE
| REAGENT or RESOURCE | SOURCE | IDENTIFIER |
| --- | --- | --- |
| **Antibodies** |  |  |
| Anti V5-Tag Antibody, clone SV5-Pk1 | Bio-Rad | MCA1360 |
| Monoclonal anti-AID/IAA17 | Cosmo Bio | CAC-APC004AM-T |
| Monoclonal anti-Rpo21, 8WG16 clone | Sigma-Aldrich | 05-952-I |
| Anti-acetylated Smc3 | (Beckouët et al., 2010) | N/A |
| **Chemicals, peptides, and recombinant proteins** |  |  |
| α-factor | Synthesized by Eurofins Genomics | N/A |
| PRONASE Protease, Streptomyces griseus | Calbiochem | 9036-06-0 |
| Benomyl | Sigma Aldrich | 17804-35-2 |
| Indole-3-acetic acid (IAA) | Sigma Aldrich | I2886 |
| Zymolyase 100T | Nacalai Tesque | 07665-55 |
| MNase | NEB | M0247S |
| **Critical commercial assays** |  |  |
| KAPA SYBR Fast qPCR kit | Sigma Aldrich | KR0389 |
| ZymoClean and Concentration Kit | ZymoResearch | D4014 |
| ZymoClean Gel DNA Recovery Kit | ZymoResearch | D4007 |
| NEBNext Ultra II DNA library Prep Kit for Illumina | NEB | E7645 |
| QIAquick PCR purification kit | QIAGEN | 28104 |
| **Deposited data** |  |  |
| ChIP-seq and Micro-C raw data | SRA | PRJNA1041821 |
| ChIP-seq and Micro-C processed data | NCBI GEO | GSE248144 |
| **Experimental models: Organisms/strains** |  |  |
| *Saccharomyces cerevisiae*, derived from BY4741 or W303-1A | See Supp.Table S1 |  |
| *Candida glabrata*, NCYC No.388 |  |  |
| **Software and algorithms** |  |  |
| Bowtie | (Langmead et al., 2009) |  |
| DROMPA | (Nakato et al., 2013) |  |
| deepTools | (Ramírez et al., 2016) |  |
| Integrative Genomics Viewer | (Robinson et al., 2011) |  |
| Juicer Tools | (Durand et al., 2016) |  |
| Galaxy HiCExplorer | (Wolff et al., 2020) |  |
| Coolpup.py | (Flyamer et al., 2020) |  |
| Cooler | (Abdennur & Mirny, 2020) |  |
| R | (R Core Team, 2021) |  |

### Yeast strains

*S. cerevisiae* strains used in this study are wild-type BY4741 and its derivatives, except the strains to visualize *URA3* locus by TetO/TetR-GFP mark, which are W303-1A derivatives. They are listed in Table S1. Epitope tagging, *aid* module tagging, and gene deletion were performed by one-step PCR-based strategy (Mathur & Kaiser, 2014; Pakchuen et al., 2016). Correct tagging or deletion of the target gene was checked by PCR. The tagging was also confirmed by western blotting.

### Cell culture

Yeast cells were cultured in YPD medium (Amberg et al., 2005) at 23°C unless otherwise mentioned. To be arrested in metaphase, cells were cultured in medium containing 80 μg/mL benomyl (Sigma-Aldrich) for 2.5 h. For synchronization in S phase, cells were first arrested in G1 phase by culturing in medium containing 2 μM α-factor (peptide synthesized by Eurofins Genomics) for 2.5 h. Then, cells were released from the arrest by transferring them to α-factor-free medium containing 150 μg/mL of Pronase (Calbiochem) and cultured for 30 min. To prepare metaphase-arrested cells lacking Scc2 protein, which were used for Micro-C analysis, *SCC2-AID* cells arrested in G1 phase by α-factor were released into the medium containing 150 μg/ml Pronase, 80 μg/mL benomyl and cultured for 1.5 h. Then, 3-indoleacetic acid (IAA) (Sigma-Aldrich, I2886) was added to the final concentration of 0.2 mg/mL and cultured for additional 2.5 h. IAA was dissolved in ethanol at 20 mg/ml. For vehicle-treated control cells, ethanol was added to the final concentration of 1% (v/v) instead of IAA. To make cells proceed through S phase without Scc2 and arrest at metaphase, *SCC2-AID* cells arrested in G1 phase by α-factor were released into the medium containing 150 μg/ml Pronase, 80 μg/mL benomyl, and 0.2 mg/mL IAA and cultured for 2.5 h. To monitor chromosome segregation in the absence of Scc2, asynchronous *SCC2-AID* cells were cultured in the medium containing 80 μg/mL benomyl for 1.5 h, followed by addition of 0.2 mg/mL IAA and cultivation for additional 2 h. The cells were, then, released from the arrest by transferring into benomyl-free medium containing 0.2 mg/mL IAA and 2 μM α-factor. To deplete Pds5 protein in metaphase-arrested cells, *PDS5-AID* cells arrested in G1 phase by α-factor were released into the medium containing 150 μg/ml of Pronase, 80 μg/mL benomyl, and 0.2 mg/mL IAA and cultured for 2.5 h. To prepare the cells that proceeded through S phase without Eco1 protein and arrested in metaphase (used for ChIP-seq analysis), we utilized *GAL-^R^ECO1* strain (Yoshimura et al., 2021), where Eco1 with short protein half-life is expressed from the galactose-dependent *GAL1* promoter. The cells grown in YPRG medium (YPD lacking glucose and supplemented with 2% raffinose and 0.1% galactose) at 30°C were transferred to YPD and cultured for 1 h at 23°C, followed by addition of 2 μM α-factor and cultivation for additional 2.5 h. Then, the cells were released into α-factor-free medium containing 150 μg/ml Pronase, and 80 μg/mL benomyl and cultured for 2.5 h. To prepare metaphase-arrested cells that lacked Eco1 after DNA replication completion (used for chromosome segregation analysis), asynchronous *GAL-^R^ECO1* cells were cultured in YPRG containing 80 μg/mL benomyl at 30°C for 1.5 h, then transferred into YPD medium containing 80 μg/mL benomyl and cultured at 23°C for 2.5 h. To observe chromosome segregation phenotype, the metaphase-arrested cells were released into benomyl-free YPD medium containing 2 μM α-factor at 23°C. The control cells that continue to express Eco1 were prepared by using YPRG medium continuously instead of YPD.

### Flow cytometry

Cell cycle synchronization was monitored by flow cytometry as described (Pakchuen et al., 2016). The analysis was conducted on a FACSCalibur or Accuri C6 flow cytometer (BD Biosciences).

### Protein analysis

Gene tagging and protein degradation were monitored by western blotting. Yeast lysate was prepared using the trichloroacetic acid (TCA) method (Pakchuen et al., 2016; Wright et al., 1989). SDS-PAGE, and Western blotting were performed as described (Blancher & Jones, 2001). Western blot image was acquired by ImageQuant LAS 4000 (GE Healthcare) or Fusion FX (Vilber). Antibodies used for ChIP and Western blotting are described in the Key Resources Table..

### Cell imaging

For fixation, 1 mL of cell culture was fixed with 3.7% formaldehyde (Fujifilm, 064-00406) for 20 min at room temperature. Then, the cells were pelleted, resuspended in 1 mL of 100% ethanol (Fujifilm, 057-00456), and incubated at 4°C for 1 hr for additional fixation.. The cells were washed twice with TBS, followed by nuclear staining in 50 µL of PBS containing 2 µg/mL DAPI (Dojindo, 340-07971) for 5 min. Finally, the cells were resuspended in 50 µL of PBS and observed by Olympus IX-83 fluorescence microscope equipped with DP23M digital camera.

### Chromatin immunoprecipitation (ChIP)

ChIP was performed as described previously (Katou et al., 2003). To conduct calibrated ChIP-seq, we utilized the *Candida glabrata* strain in which the endogenous Scc1 gene was tagged with PK epitopes (a gift from Prof. Kim Nasmyth). Every 2 (or 5 in the case of Scc2 ChIP-seq) OD units of *S. cerevisiae* cell culture were mixed with 1 OD unit of *C. glabrata* asynchronous cell culture immediately before fixation.

### ChIP-seq library preparation and sequencing

Both input and ChIP fractions of DNA were processed. Purified DNA was further fragmented to the size of ∼150 bp by the Covaris focused-ultrasonicator (Covaris M220) and subjected to library preparation using the NEBNext Ultra II DNA Library Prep Kit for Illumina (NEB, E7645). Sequencing of the library was performed using Illumina HiSeq2500 and NextSeq2000 to produce 65-bp single-end reads and 36-bp paired-end reads, respectively.

### Calibrated ChIP-seq data analysis

Raw sequencing reads were mapped to *S. cerevisiae* and *C. glabrata* reference genomes by Bowtie (Langmead et al., 2009) using default parameters. For a specific ChIP sample, the number of non-redundant reads mapped to *S. cerevisiae* and *C. glabrata* genome in the input and ChIP fractions were used to calculate the Occupancy Ratio (ORs) by the following formula as described previously (Hu et al., 2015).

OR = (ChIP_Scer_ × input_Cgla_)/(input_Scer_ × ChIP_Cgla_)

where ChIP_Scer_, the number of the reads that were in the ChIP sample and mapped to *S. cerevisiae* genome;

input_Scer_, the number of the reads that were in the input sample and mapped to *S. cerevisiae* genome;

ChIP_Cgla_, the number of the reads that were in the ChIP sample and mapped to *C. glabrata* genome;

input_Cgla_, the number of the reads that were in the input sample and mapped to *C. glabrata* genome.

Then, to enable quantitative comparison among multiple ChIP experiments, the normalization factor (NF) of each sample was calculated as described in (Orlando et al., 2014).

NF = OR_treated_/OR_control_

where OR_treated_, the OR for the target condition;

OR_treated_, the OR for the reference condition.

The non-redundant reads mapped to *S. cerevisiae* genome were subject to binning using parse2wig in the DROMPAplus package (version 1.17.2) (Nakato et al., 2013), and the results was output as a file in wig format. For a ChIP sample, the following normalization option was applied; ‘-n GR –nrpm X’, where X is the product of 1,500,000 and NF for the experiment. Finally, the wig files for input and ChIP samples of a specific ChIP experiment were fed to PC_ENRICH function in DROMPAplus to generate a file showing normalized ChIP/Input fold-enrichment ratio for each 100-bp genome bin in bigwig or bedgraph format, which was used for downstream analyses. Aggregation plot and heatmap were produced using deepTools (Ramírez et al., 2016). To visualize ChIP-seq profiles, Integrative Genomics Viewer (Robinson et al., 2011) was used. The ChIP-seq peak regions, where the ChIP/Input fold-enrichment ratio was equal to or above the threshold of 2.0, were identified by DROMPA’s PC_ENRICH function. The statistics of ChIP-seq sequencing are in Supplement Table S3.

### ChIP-qPCR

Analysis of ChIP-purified DNA by quantitative PCR (qPCR) was performed using KAPA SYBR Fast qPCR kit (KAPA Biosystems) and StepOnePlus real-time PCR systems (Life Technology, Inc.) as manufacturers’ instructions. Input and ChIP DNA were measured in duplicate. The primers used in qPCR are listed in Table S2.

### Micro-C library preparation

Micro-C was conducted as previously described (Costantino et al., 2020; Hsieh et al., 2016) with minor modifications. Briefly, 100 mL of culture at OD ∼ 0.8 was fixed with 3% formaldehyde for 15 min at 30^°^C. Fixation was quenched by adding glycine to a final concentration of 0.2 M and shaking for 5 min at 30^°^C. Cells were collected and washed with sterile, ice-cold water before processing to cell-wall permeabilization by resuspending to 10 mL buffer Z (1 M sorbitol, 50 mM Tris-HCl pH 7.4, supplemented with 10 mM 2-Mercaptoethanol (Wako, 133-14571)) containing 250 μg/mL Zymolyase 100T (Nacalai Tesque, 07665-55), and shaking at 180 rpm for 40 min at 30^°^C. The spheroplasts were collected and washed by PBS, then transferred to a 1-mL DNA LoBind tube (Eppendorf, 022431021). Cells were resuspended to 1 mL PBS, and 10 μL of 0.3 M DSG (Thermo Fisher Scientific, A35392) was added to the final concentration of 3 mM for the second cross-linking. The tube was shaken for 40 min at 30^°^C before quenching by 0.2 M glycine for 5 min at 30^°^C on a shaking incubator. Cells were washed by PBS and pelleted by centrifugation. The cell pellets can be snap-frozen in liquid nitrogen and stored at-80°C or immediately processed to the next step.

For chromatin fragmentation by MNase, cell pellets were first incubated for 5 min in 200 μL ice-cold MB1 (50 mM NaCl, 10 mM Tris-HCl pH 7.5, 5 mM MgCl_2_, 1 mM CaCl_2_, 0.075% NP-40 and 1× cOmplete EDTA-free protease inhibitor cocktail (Roche, 04693132001)). ∼1,000 U of MNase (NEB, M0247S) was added to every cell derived from 100-mL cell culture, and the mixture was incubated for 20 min at 37^°^C, which produced ∼90% mono-nucleosome and ∼10% di-nucleosome. The reaction was stopped by adding 5 μL of 0.1M EGTA (final conc. 2.5 mM), incubated at 65°C for 10min. In-nucleus fragmented chromatin was pelleted and washed twice by MB2 (50 mM NaCl, 10 mM Tris-HCl pH 7.5, 10 mM MgCl_2_) before processing to end-preparation.

A 3-step end-preparation was conducted according to (Costantino et al., 2020). The reaction was inactivated by adding EDTA to the final concentration of 30 mM and incubating for 20 min at 65°C. Chromatin was washed once by 1× T4 DNA ligase buffer (NEB, B0202) and pelleted by centrifugation.

For proximity ligation, the pellet was resuspended into 500 μL of ligation mix (1× T4 DNA ligase buffer, 1× BSA (NEB, B9000S), 5000 CEU T4 DNA ligase (NEB, M0202L)) and rotated on a slow rotator for 3 hours or more at room temperature. The biotin-dNTP at unligated ends was then removed by incubation with 200 U Exonuclease III (NEB, M0206S) in 1× NEB buffer 1 (NEB, B7001S) for 15 min at 37°C. In the next step, chromatin was processed to reverse crosslinking by incubating at 65°C overnight in TE/1%SDS supplemented with 15 μg/mL Proteinase K and 15 μg/mL RNase A. DNA was purified using standard phenol:chloroform extraction and ethanol precipitation methods, and was resuspended in 100 uL elution buffer (buffer EB, QIAgen, 19086). DNA was further purified and concentrated by Zymo Clean and Concentrator Kit (ZymoResearch, D4014) and eluted to 15μL buffer EB. To select DNA fragments ranging from 250 to 350bp in size, DNA samples were loaded on 3% Nusieve^TM^ GTG^TM^ Agarose gel (Lonza, 50081), and electrophoresis was run at 100V in 1X TBE buffer for ∼50 min. The DNA corresponding to the desired size was excised and purified with ZymoClean Gel DNA Recovery Kit (ZymoResearch, D4007) and finally eluted to 50 µL by buffer EB. To pull down the biotin-labeled DNA, the samples were incubated with 5 µL of Dynabeads MyOne Streptavidin C1 beads (Invitrogen, 65001) on a slow rotator at room temperature for 20 min. The DNA-bound streptavidin beads were washed twice with 200 µL TWB (5 mM Tris-HCl pH 7.5, 1 mM EDTA, 1 M NaCl, 0.05% Tween 20) and once with 200 µL buffer EB, followed by on-beads library preparation using the NEBNext Ultra II DNA Library Prep Kit for Illumina according to the manufacturer instruction from the end repair to adaptor ligation step. After adapter ligation step, beads were washed twice with 200 µL TWB and once with 200 µL buffer EB, resuspended into 15 µL buffer EB, then processed to PCR amplification step following the protocol from NEBNext Ultra II DNA Library Prep Kit for Illumina. Finally, the DNA library was cleaned twice using AMPure XP Reagent (Beckman Coulter, A63881) and eluted to 25 µL by buffer EB. Libraries were sequenced by HiSeqX to generate ∼100 million of 150-pb paired-end reads per sample.

## Micro-C data analysis

### • Contact matrix generation

Sequencing reads were processed to generate a visualizable contact matrix, following the previously described analysis pipeline (*Micro-C Documentation Release 0.1 Dovetail*, 2023), which utilizes bwa for mapping (Li, 2013) and pairtools (Abdennur et al., 2023) for processing sequencing reads. For comparison between samples, the valid read pairs of different samples were normalized to the same total read number before generating contact matrices. Juicer Tools (Durand et al., 2016) was used to convert.pairs file into a.hic contact matrix that stored the mapped pairs in 5 resolutions (100bp, 200bp, 500bp, 1kb, and 2kp) and 4 normalizations (VC, VC_SQRT, KR, and SCALE), which is ready to be visualized by the compatible web-based software Juicebox (https://www.aidenlab.org/juicebox/). The statistics of Micro-C sequencing is in Supplement Table S4.

### **•** Count-versus-distance decaying curve and its first derivative

Iterative correction (Imakaev et al., 2012) was applied to the contact matrix at the resolution of 500 bp using HiCExplorer’s hicCorrect module.The resultant IC-corrected matrix was subjected to HiCExplorer’s hicPlotDistVsCount module (Wolff et al., 2020) to calculate and produce a result matrix of the number of reads at every 500-bp bin size distance. The output matrix was then used to plot the log scale decaying curve and its first derivative by R.

### **•** Loop calling

For loop calling,.hic files were subjected to Juicer HICCUPS algorithm (Durand et al., 2016) with the following parameters: “-k KR-r 1000-f 0.1-p 8-i 16-d 2500”, which scanned the contact matrix (normalized by Knight-Ruiz (KR) balancing (Knight & Ruiz, 2013) and stored in the.hic files at 1kb resolution) using a 16×16-pixel window to search for enriched pixels of peak-width 8. The enriched pixels separated by a distance less than 2500 bp were merged and then filtered by a false discovery rate of 0.1 to generate the final loops list in a bedpe-format file.

### **•** Aggregate peak analysis

The aggregate peak analysis (APA) to present the average contact enrichment score around a set of genome loci (on-diagonal APA) or between pairs of specific loci (off-diagonal APA) was conducted using the Coolpup.py package (Flyamer et al., 2020). To standardize the contact level with the background, we first calculated the expected interactions for each chromosome arm and then computed the observed-over-expected (observed/expected) enrichment score. The 200-bp resolution contact matrix in.cool format was balanced to filter out poorly mapped bins by *cooler balance* (Abdennur & Mirny, 2020). The expected interaction was calculated from the balanced contact matrix using the *expected_cis* API module of the cooltools package (Abdennur et al., 2022). We, then, fed the following three inputs into *Coolpup.py* to calculate and plot the average observed/expected score between pairs of specific sites: the balanced contact matrix, the expected interaction matrix, and the list of the genome loci or paired loci to be analyzed in bedpe format. The list of paired loci of which the averaged pile-up was computed was either a loop file produced by HICCUPS (Durand et al., 2016), or a manually generated bedpe file containing, for example, a collection of pairs between a centromeric anchor and a cohesin-bound anchor on chromosome arms grouped by distances between two anchors (used in Figure 5B). *coolpup.py* was executed with the parameters “–flank 10000 –mindist 5000”, which will calculate the average observed/expected score of interactions between the designated pairs of loci separated by a distance larger than 5 kb, in a 20×20-kb window centered at the enriched pixels representing the interaction between two loci. This execution generated a matrix data, which was subsequently used to plot the pile-up heatmap by *plotpup.py* command.

### **•** Insulation score calculation and chromosomal domain analysis

The KR-balanced contact matrix at the resolution of 1 kb was subjected to the hicFindTADs module of the HiCExplorer (Wolff et al., 2020). The parameters used were “--minDepth 5000--maxDepth 10000--step 5000--correctForMultipleTesting fdr--thresholdComparisons 0.01--delta 0.01”, which will compute the insulation score on the 1 kb contact matrix at window sizes of 5000 and 10000 using the false discovery rate (FDR) for multiple comparisons with the q-value threshold set to 0.01. The output files consist of lists of boundaries and domains in bed format and genome-wide insulation score in bedGraph format, which can be used to plot the average profile.

## Data availability

All the sequencing data for ChIP-seq and Micro-C are available through the GEO accession number GSE248144.

