## Supplemental Figures & Tables for "Controlling mechanism of the Scc2-cohesin interaction to restrict peri-centromeric DNA loop expansion and facilitate mitotic chromosome segregation"

Figure S1

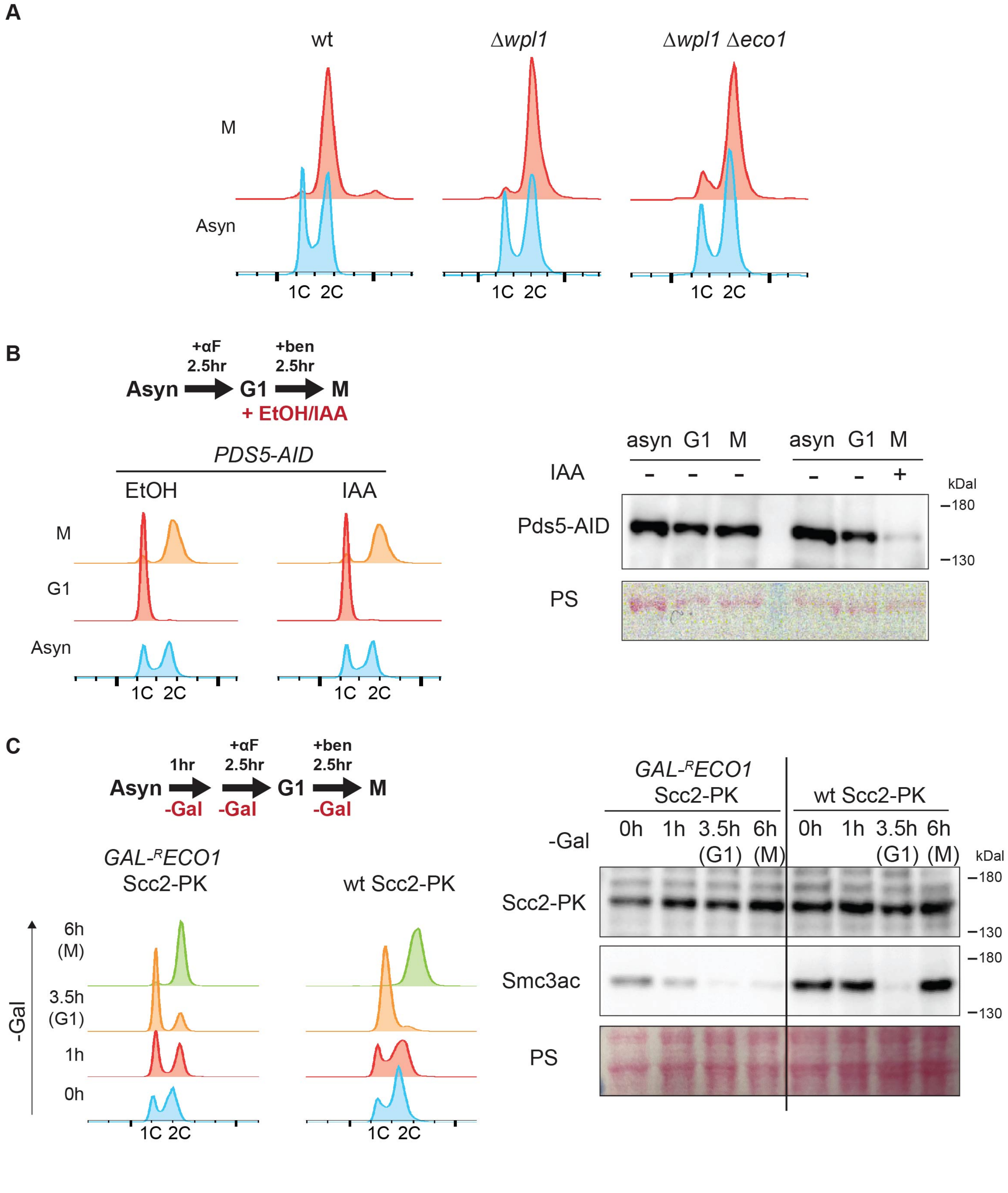
A
wt
Δwpl1
Δwpl1 Δeco1
M
Asyn
1C 2C
B
+αF
2.5hr
+ben
2.5hr
Asyn
G1
M
+ EtOH/IAA
PDS5-AID
EtOH
IAA
G1
1C 2C
asyn G1 M asyn G1 M
IAA
- - - - - +
kDal
Pds5-AID
180
130
PS
C
1hr
+αF
2.5hr
+ben
2.5hr
-Gal
GAL-RECO1
Scc2-PK
wt Scc2-PK
6h
(M)
3.5h
(G1)
1h
0h
0h 1h 3.5h (G1) 6h (M)
Smc3ac

**Figure S1. Validation of cell cycle arrest and protein depletion**
**(A)** Flow cytometry data to monitor cell cycle arrest in wt, Δ*wpl1*, and Δ*wpl1* Δ*eco1*. Asyn, asynchronous; M, metaphase.
**(B)** (Left) Cell culture condition and flow cytometry data to monitor cell cycle arrest for *PDS5-AID.* To deplete Pds5 protein in metaphase-arrested cells, *PDS5-AID* cells arrested in G1 phase by α-factor (αF) were released into the medium containing benomyl (Ben) and idole-3-acetic acid (IAA) and cultured for 2.5 h. (Right) Western blot to verify IAA-dependent degradation of Pds5-AID. PS, Ponceau S staining as loading control.
**(C)** (Left) Cell culture condition and flow cytometry data to monitor cell cycle arrest for *GAL-$^{R}$ECO1*. To prepare the cells that proceeded through S phase without Eco1 protein and arrested in metaphase, *GAL-$^{R}$ECO1* cells grown in galactose-containing medium were transferred to galactose-free medium (YPD) and cultured for 1 h at 23°C, followed by addition of 2 μM α-factor and cultivation for additional 2.5 h. Then, the cells were released into α-factor-free YPD containing benomyl and cultured for 2.5 h. Gal, galactose. (Right) Western blot to verify the depletion of Eco1 in Gal-free medium. Eco1 depletion resulted in the disappearance of acetylated cohesin (Smc3ac) in metaphase.

**Figure S2**

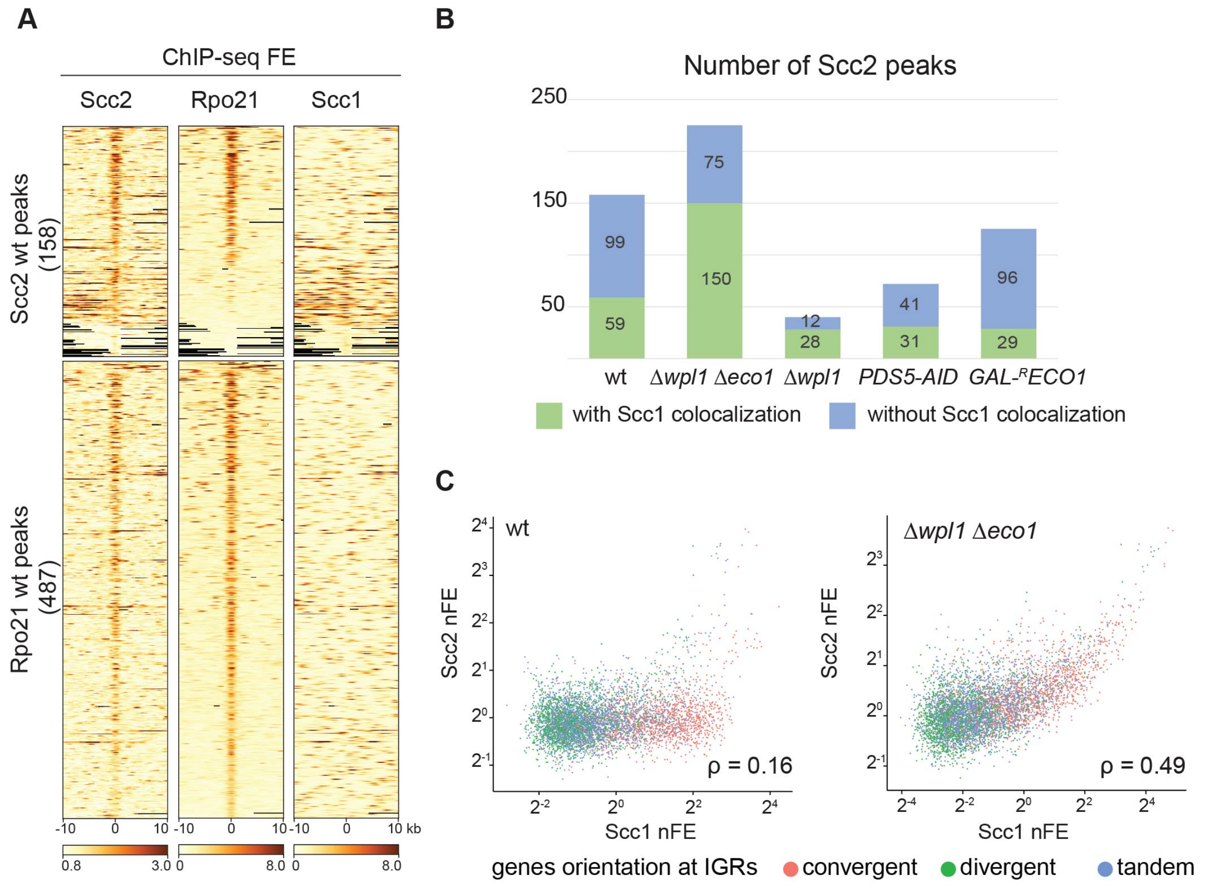


**Figure S2. Colocalization of Scc2 with RNA pol II in wt and with cohesin in Δ*wpl1* Δ*eco1***
**(A)** Heatmap of Scc2, Rpo21 (the largest subunit of RNA pol II), and Scc1 ChIP-seq FE in wt. 10-kb surrounding regions of Scc2 and Rpo21 peaks in wt are depicted. Regions are sorted in descending order of Rpo21 FE in wt.
**(B)** The number of Scc2 peaks in the genome of the indicated strains. Green, those overlapping with the Scc1 peaks; blue, those not overlapping with the Scc1 peaks.
**(C)** Correlation between Scc1 and Scc2 ChIP-seq nFE in wt and Δ*wpl1* Δ*eco1*. Each dot corresponds to an intergenic region (IGR). The color of the dot indicates the orientation of the genes adjacent to the IGR. ρ, Spearman's correlation coefficient.

**Figure S3**

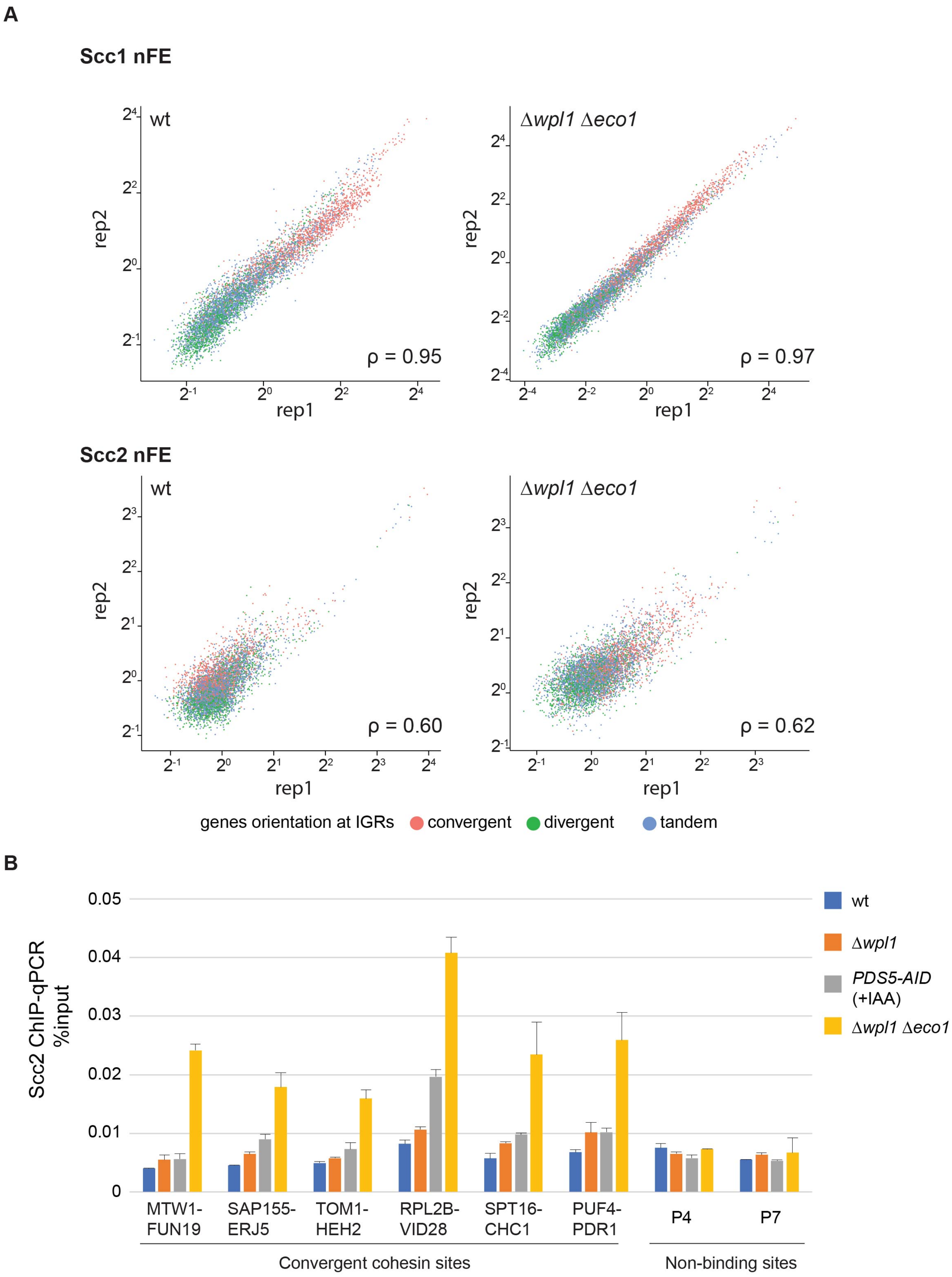
A
Scc1 nFE
wt
Δwpl1 Δeco1
rep2
rep1
ρ = 0.95
ρ = 0.97
Scc2 nFE
ρ = 0.60
ρ = 0.62
genes orientation at IGRs
convergent
divergent
tandem
B
Scc2 ChIP-qPCR %input
0.05
0.04
0.03
0.02
0.01
0
MTW1-FUN19
SAP155-ERJ5
TOM1-HEH2
RPL2B-VID28
SPT16-CHC1
PUF4-PDR1
P4
P7
Convergent cohesin sites
Non-binding sites
wt
Δwpl1
PDS5-AID (+IAA)
Δwpl1 Δeco1

**Figure S3. Reproducibility and validation of ChIP-seq data**

**(A)** Correlation between two biological replicates of Scc1 or Scc2 ChIP-seq. nFE values at all intergenic regions (IGRs) are depicted. Results in wt and Δ*wpl1* Δ*eco1* are shown. The color of the dot indicates the orientation of the genes adjacent to the IGR. ρ, Spearman's correlation coefficient.

**(B)** ChIP-qPCR of Scc2 in wt, Δ*wpl1*, *PDS5-AID* (+IAA) and Δ*wpl1* Δ*eco1*. Analyzed loci are six representative cohesin binding sites and two negative control sites (Non-binding sites). See table S2 for detailed information on the primers used. The mean of two biological replicates was shown. Error bar, standard deviation.

### Figure S4

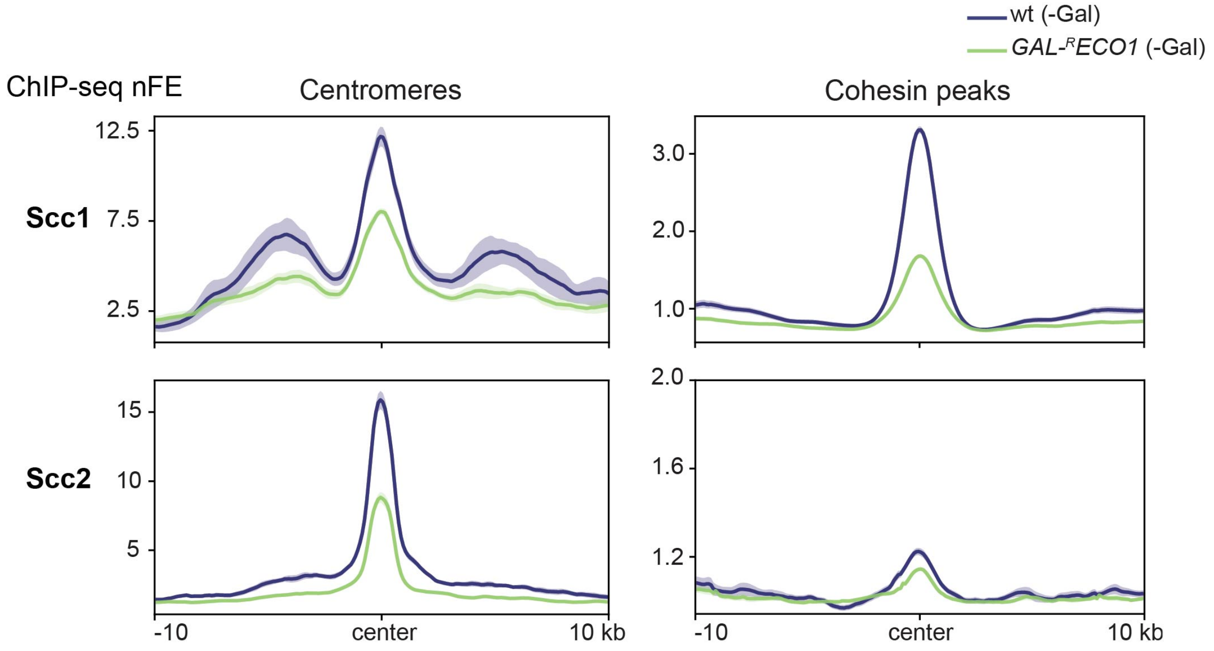


**Figure S4. Scc1 and Scc2 chromosomal binding in *GAL-$^{R}$ECO1***

Aggregated ChIP-seq profiles of Scc1 and Scc2 in wt and *GAL-$^{R}$ECO1* cultured in galactose-free medium. The profiles are centered at the centromeres or the summit of non-centromeric Scc1 peaks in wt. Bold line, mean; shaded area, 95% confidence interval.

**Figure S5**

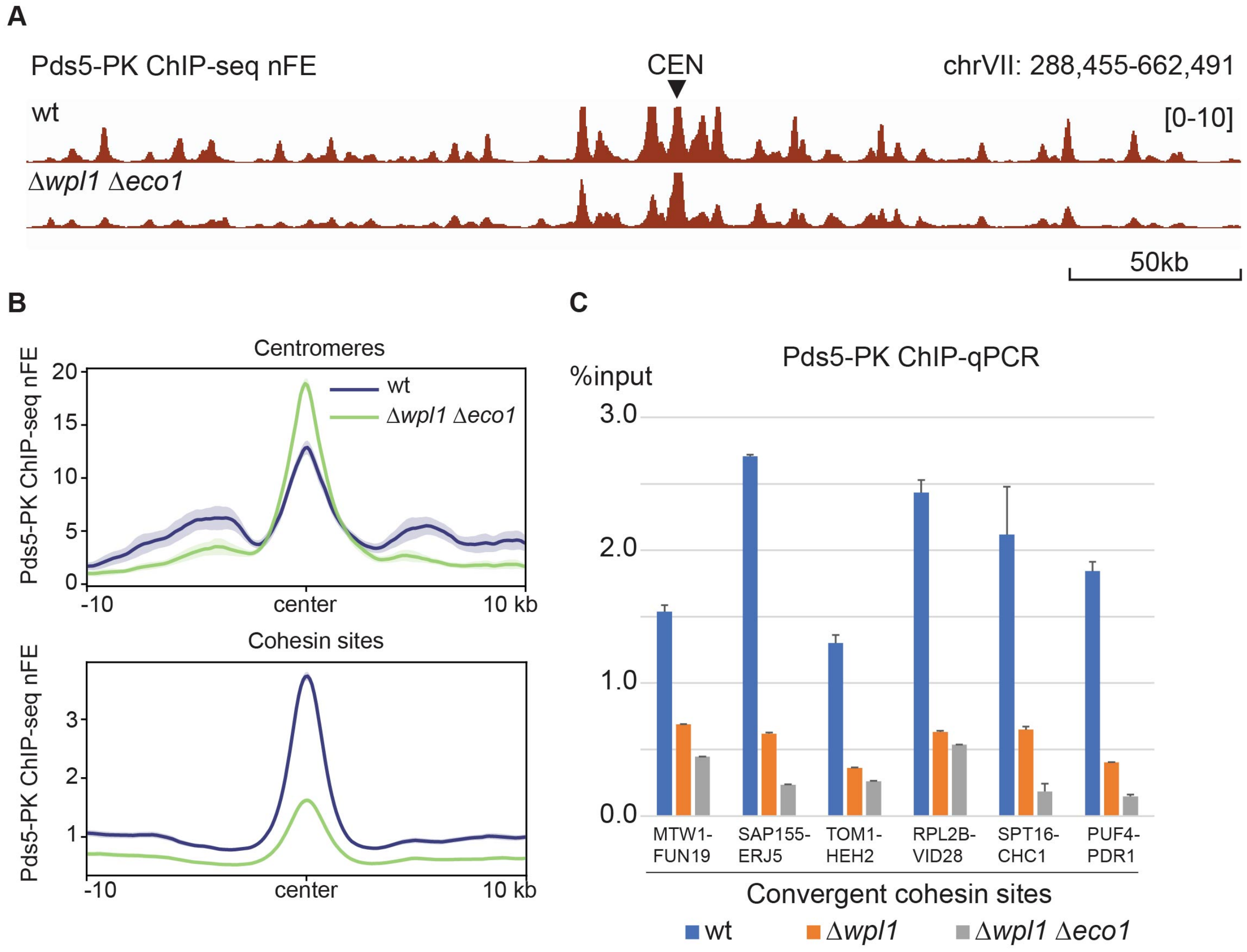


**Figure S5. Impaired Pds5 co-localization at the cohesin sites in Δ*wpl1* Δ*eco1***

**(A)** Calibrated ChIP-seq profiles of Pds5-PK in wild-type and Δ*wpl1* Δ*eco1*. Cells are arrested at metaphase by benomyl treatment.

**(B)** Aggregated ChIP-seq profiles of Pds5 in wt and Δ*wpl1* Δ*eco1*. The profiles are centered at the centromeres or the summit of non-centromeric Scc1 peaks in wt. Bold line, mean; shaded area, 95% confidence interval.

**(C)** ChIP-qPCR of Pds5 in wt, Δ*wpl1,* and Δ*wpl1* Δ*eco1*. Analyzed loci are six representative cohesin binding sites. The mean of two technical replications was shown. Error bar, standard deviation.

**Figure S6**

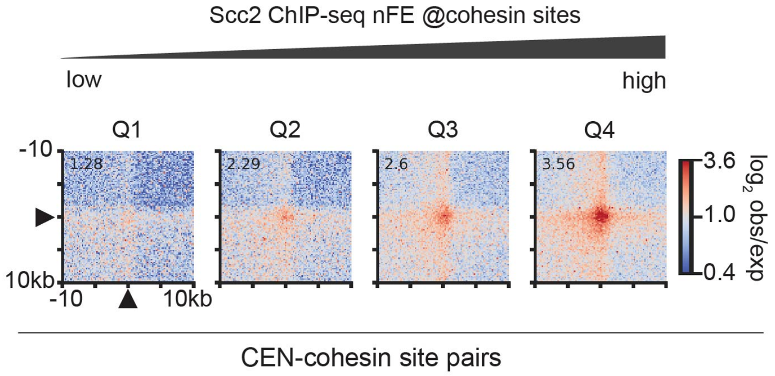


**Figure S6. Correlation between Scc2 chromosomal binding and contact frequency of the pericentromeric DNA loops**

Average contact frequency (represented as observed/expected ratio) between a centromere and a cohesin site on the same chromosome in Δ*wpl1* Δ*eco1*. The centromere-cohesin site pairs were divided into four groups according to the Scc2 nFE value at the cohesin site (Q1 to Q4 in order of increasing FE), as described in Figure 1C. Black triangles indicate the two anchor loci of each pair. The number in the top-left corner of each plot indicates the average enrichment score of the 3 x 3 central pixels.

Figure S7

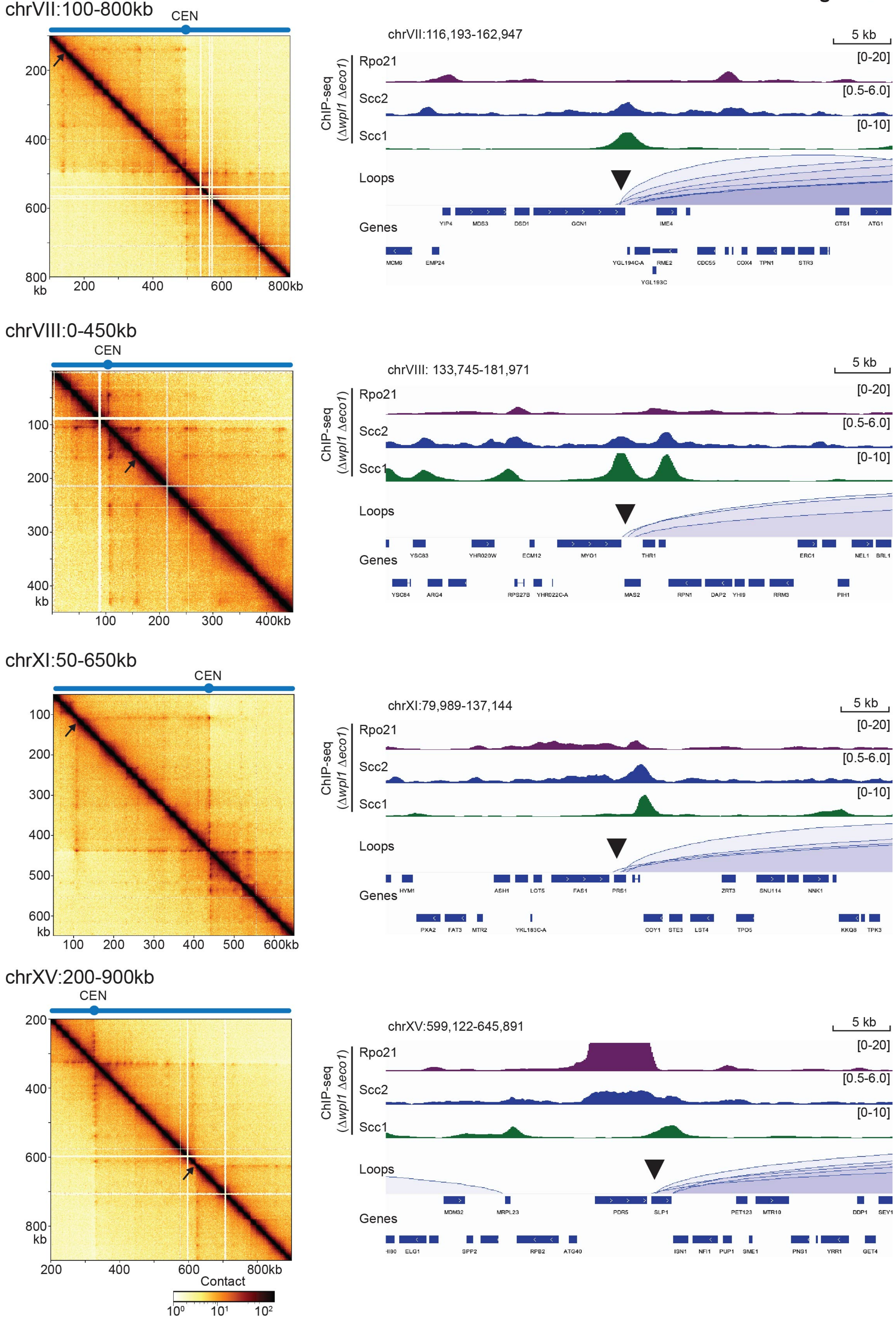
chrVII:100-800kb
CEN
200
400
600
800
kb
200
400
600
800kb
chrVII:116,193-162,947
5 kb
ChIP-seq
(Δwpl1 Δeco1)
Rpo21
[0-20]
Scc2
[0.5-6.0]
Scc1
[0-10]
Loops
Genes
chrVIII:0-450kb
CEN
100
200
300
400
kb
100
200
300
400kb
chrVIII: 133,745-181,971
5 kb
ChIP-seq
(Δwpl1 Δeco1)
Rpo21
[0-20]
Scc2
[0.5-6.0]
Scc1
[0-10]
Loops
Genes
chrXI:50-650kb
CEN
100
200
300
400
500
600
kb
100
200
300
400
500
600kb
chrXI:79,989-137,144
5 kb
ChIP-seq
(Δwpl1 Δeco1)
Rpo21
[0-20]
Scc2
[0.5-6.0]
Scc1
[0-10]
Loops
Genes
chrXV:200-900kb
CEN
200
400
600
800
kb
200
400
600
800kb
Contact
$10^0$
$10^1$
$10^2$
chrXV:599,122-645,891
5 kb
ChIP-seq
(Δwpl1 Δeco1)
Rpo21
[0-20]
Scc2
[0.5-6.0]
Scc1
[0-10]
Loops
Genes

**Figure S7. Potential strong barrier sites for loop extrusion found in chromosome arms of Δ*wpl1* Δ*eco1***
(Left) Micro-C contact map at 1-kb resolution in the indicated genome regions. Potential strong barrier sites for loop extrusion are marked with black arrows. CEN, centromere. (Right) ChIP-seq profiles of Rpo21, Scc2 and Scc1 in Δ*wpl1* Δ*eco1* around the potential barrier sites (black arrowheads). The detected DNA loops (Loops), as well as the genes annotated in the RefSeq (Genes), are also shown.

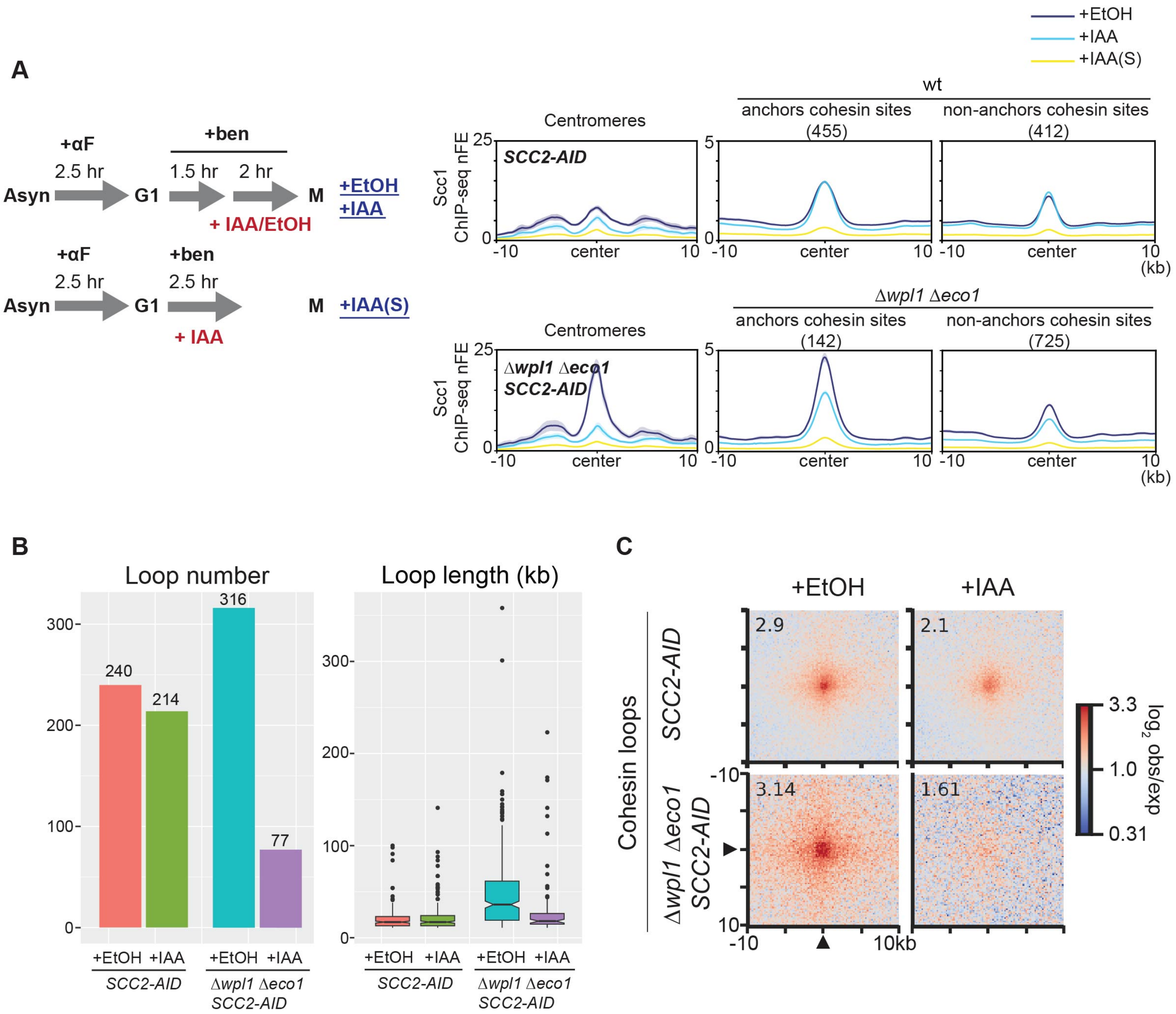


**Figure S8. Validation of culture condition and features of DNA loops in Scc2-depleted cells**

**(A)** (Left) Schematic representation of the experimental protocol used to arrest cells in metaphase and induce rapid degradation of Scc2-AID. The cells were arrested in metaphase before Scc2 depletion in the '+IAA' condition while the cells proceeded through S phase without Scc2 in the '+IAA(S)' condition. (Right) Aggregated plots of Scc1 ChIP-seq nFE in *SCC2-AID* and Δ*wpl1* Δ*eco1* *SCC2-AID* strains cultured under the indicated conditions (+EtOH, +IAA, +IAA(S)). 10-kb surrounding regions of the centromeres, anchor cohesin sites, and non-anchor cohesin sites are averaged and depicted. Bold line, mean; shaded area, 95% confidence interval.

**(B)** Comparison of the number and length of the DNA loops detected in the indicated samples.

**(C)** Average contact frequency of the cohesin loops (loops connecting between cohesin binding sites) detected in vehicle-treated *SCC2-AID* and Δ*wpl1* Δ*eco1* *SCC2-AID* (+EtOH). The averaged contact frequency for the same locus-pair in IAA-treated condition was shown side by side for comparison (+IAA).

Figure S9

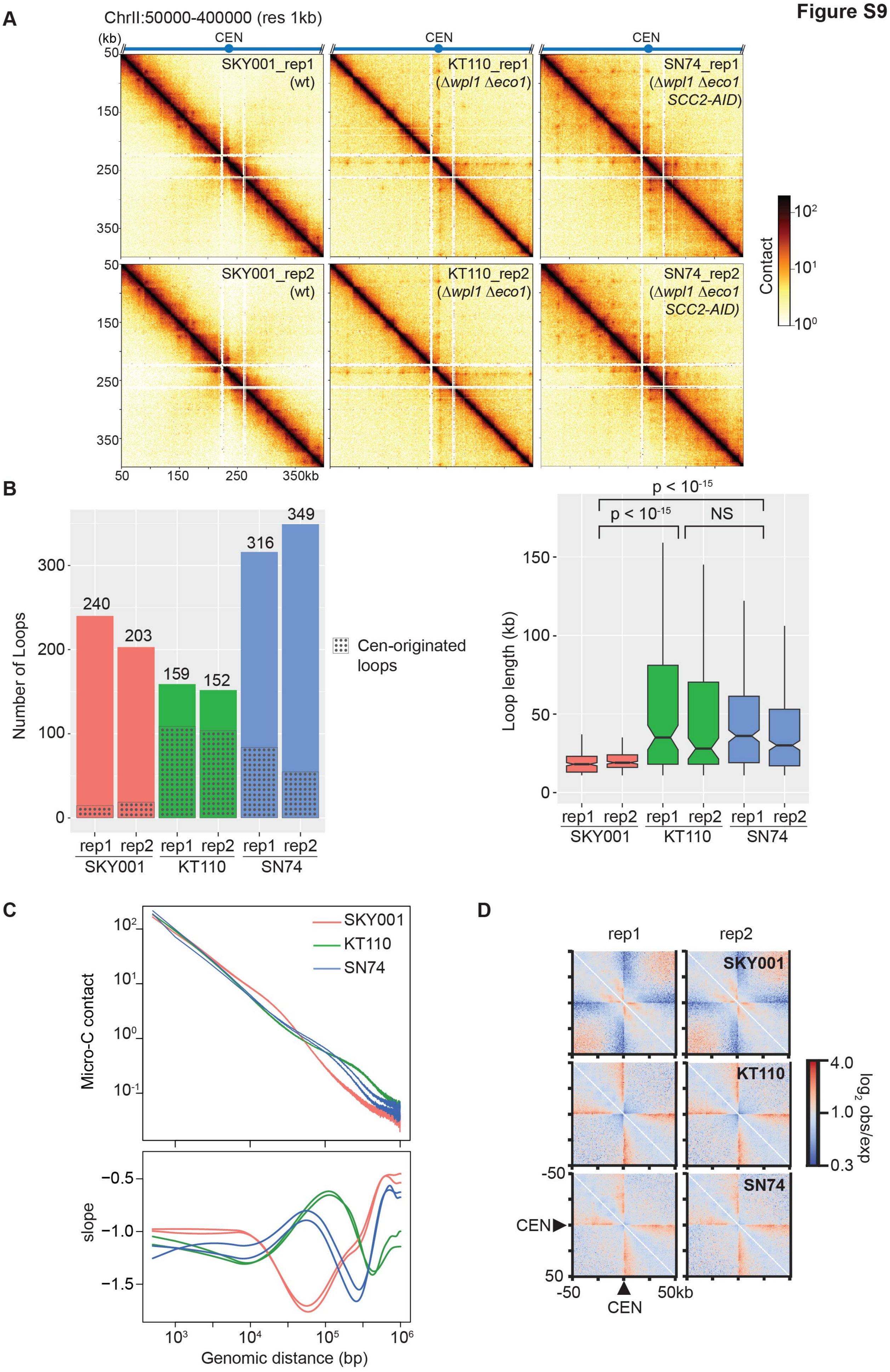
A
ChrII:50000-400000 (res 1kb)
(kb)
CEN
SKY001_rep1 (wt)
KT110_rep1 (Δwpl1 Δeco1)
SN74_rep1 (Δwpl1 Δeco1 SCC2-AID)
SKY001_rep2 (wt)
KT110_rep2 (Δwpl1 Δeco1)
SN74_rep2 (Δwpl1 Δeco1 SCC2-AID)
50
150
250
350
350kb
Contact
B
Number of Loops
240
203
159
152
316
349
Cen-originated loops
rep1 rep2 rep1 rep2 rep1 rep2
SKY001 KT110 SN74
p < 10-15
p < 10-15
NS
Loop length (kb)
C
SKY001
KT110
SN74
Micro-C contact
slope
Genomic distance (bp)
D
rep1
rep2
SKY001
KT110
SN74
CEN
-50
50kb
4.0
1.0
0.3
log2 obs/exp

**Figure S9. Micro-C data reproducibility and minor difference between strains**

**(A)** Contact matrix at the resolution of 1 kb of SKY001 (wt), KT110 (Δ*wpl1* Δ*eco1*), and SN74 (Δ*wpl1* Δ*eco1* *SCC2-AID SCC1-9PK*). For each strain, two biological replicates are shown. The number of valid reads in each sample was normalized to 42 million.

**(B)** The number and length of pericentromeric loops detected in the indicated samples. The outliers were not depicted in the box plot. Data from the two replicates were merged and compared between the strains. The numbers above the plots are p-values (Mann-Whitney U test, two-tailed). NS, nonsignificant difference ($p > 0.05$).

**(C)** Contact-versus-distance decaying curves of IC-corrected contact matrix and their first derivatives (slope), at the resolution of 500 bp. Results of two biological replicates for SKY001, KT110, and SN74 strains are shown.

**(D)** Average contact frequency around the centromeres in the indicated samples. Black triangles indicate the centromere position.

**Supplement Table S1. S. cerevisiae strains used in this study**

| Relative figure | Strain ID | Genotype |
|---|---|---|
| Fig.3, Fig.S9 | SKY001[a] | *MATa his3Δ1 leu2Δ0 met15Δ0 ura3Δ0 trp1Δ* |
| Fig.5 | KT127[b] | *MATa his3Δ1 leu2Δ0 met15Δ0 ura3Δ0 trp1Δ wpl1Δ::LEU2* |
| Fig.3, Fig.S7, Fig.S9 | KT110[b] | *MATa his3Δ1 leu2Δ0 met15Δ0 ura3Δ0 trp1Δ wpl1Δ::LEU2 eco1Δ::TRP1* |
| Fig.1, Fig.2, Fig.4, Fig.S1, Fig.S2, Fig.S4 | SN40 | *MATa his3Δ1 leu2Δ0 met15Δ0 ura3Δ0 trp1Δ SCC2-9PK:TRP1* |
| Fig.1, Fig.2, Fig.4, Fig.5, Fig.S1, Fig.S2 | SN54 | *MATa his3Δ1 leu2Δ0 met15Δ0 ura3Δ0 trp1Δ wpl1Δ::LEU2 eco1Δ::TRP1 SCC2-9PK:HIS3MX6* |
| Fig.1 | SN53 | *MATa his3Δ1 leu2Δ0 met15Δ0 ura3Δ0 trp1Δ wpl1Δ::LEU2 SCC2-9PK:HIS3MX6* |
| Fig.1, Fig.S4 | SN41 | *MATa his3Δ1 leu2Δ0 met15Δ0 ura3Δ0 trp1Δ SCC2-9PK:TRP1 pGAL-Ub-$^{R}$ECO1:kanMX6* |
| Fig.S4 | SN27 | *MATa his3Δ1 leu2Δ0 met15Δ0 ura3Δ0 trp1Δ SCC1-9PK:kanMX6* |
| Fig.S4 | SN39 | *MATa his3Δ1 leu2Δ0 met15Δ0 ura3Δ0 trp1Δ SCC1-9PK:TRP1 pGAL-Ub-$^{R}$ECO1:kanMX6* |
| Fig.1, Fig.5 | SN80 | *MATa his3Δ1 leu2Δ0 met15Δ0 ura3Δ0 trp1Δ SCC2-9PK:TRP1 AUR1:TIR1-9Myc,AUR1-C PDS5-AID:kanMX6* |
| Fig.S5 | SN47 | *MATa his3Δ1 leu2Δ0 met15Δ0 ura3Δ0 trp1Δ PDS5-9PK:HIS3MX6* |
| Fig.S5 | SN48 | *MATa his3Δ1 leu2Δ0 met15Δ0 ura3Δ0 trp1Δ wpl1Δ::LEU2 eco1Δ::TRP1 PDS5-9PK:HIS3MX6* |
| Fig.1, Fig.4, Fig.6, Fig.7, Fig.S1, Fig.S2, Fig.S8, Fig.S9 | SN74 | *MATa his3Δ1 leu2Δ0 met15Δ0 ura3Δ0 trp1Δ wpl1Δ::LEU2 eco1Δ::TRP1 AUR1:TIR1-9Myc,AUR1-C SCC2-AID:kanMX6 SCC1-9PK:HIS3MX6* |
| Fig.1, Fig.2, Fig.4, Fig.6, Fig.7, Fig.S1, Fig.S2, Fig.S8 | SN75 | *MATa his3Δ1 leu2Δ0 met15Δ0 ura3Δ0 trp1Δ AUR1:TIR1-9Myc,AUR1-C SCC2-AID:kanMX6 SCC1-9PK:HIS3MX6* |
| Fig.7 | ST258[b] | *MATa ade2-1 can1-100 leu2-3,112 trp1-1 ura3::URA3,tetOs his3::HIS3,tetR-GFP wpl1Δ::LEU2* (W303-1A derivative) |
| Fig.7 | ST722 | *MATa ade2-1 can1-100 leu2-3,112 trp1-1 ura3::URA3,tetOs his3::HIS3,tetR-GFP wpl1Δ::LEU2 pGAL1-Ub-$^{R}$ECO1:kanMX6* (W303-1A derivative) |

*C. glabrata* strain used for calibration (*SCC1-PK:natMX6*) is a gift from Prof. Kim Nasmyth.

[a] described in (Katou et al., 2003)
[b] described in (Sutani et al., 2009)

**Supplement Table S2. Primer pairs used for ChIP-qPCR**

| ID | Forward | Reverse | Target |
|---|---|---|---|
| **P4** | CACCGTGCTCCAAATGGCCT | AGCTTCGCTGCTTGATGCGT | chrII: 541262-541355 |
| **P7** | GCGGCCGAACGGTCACTAGA | TGAGCCAATGCTTCCAGAGGCTA | chrII: 569861-569922 |
| **P11** | AGGAACGTCCTTCGAATCCCTGA | ACGCTAACTCCACGTCGTTCACT | chrI: 79982-80107 (MTW1-FUN19) |
| **P12** | AGTCGACTTGGATGCCGCAA | AGGGTCTTCAGGACGAGATTTCAA | chrVI: 216353-216624 (SAP155-ERJ5) |
| **P15** | AAACCATATTGAAACAGCGAATGGT | TGGAGAGATCATGACATGTTGGG | chrVI: 1379871-1380069 (TOM1-HEH2) |
| **P16** | ACTACCTTTTGCAGCCGGGGT | TCTGGAACCAGGGCATTCACCA | chrIX: 318368-318569 (RPL2B-VID28) |
| **P18** | TCCACGTCTTAAAATCCTGTGGGT | CCGAAGAACTCGCTAAAAAGTCCG | chrVII: 102535-102623 (SPT16-CHC1) |
| **P20** | AGAGAAGGAGATGCCCTAGAAAACA | AGTATCCTGTGGAGCGACGTT | chrVII: 468980-469107 (PUF4-PDR1) |

**Supplement Table S3. ChIP-seq data**

| Sample | Total mapped | S.cer | | C.gla | | OR | NF | Number of normalized reads | Relative figure |
|---|---|---|---|---|---|---|---|---|---|
| | | Uniquely mapped | % | Uniquely mapped | % | | | | |
| Scc2-PK wt IP_rep1 | 6472695 | 2452200 | 37.88530125 | 2954642 | 45.64778659 | 0.097863022 | 1 | 1500000 | Fig.S2 |
| Scc2-PK wt WCE_rep1 | 7144449 | 5415436 | 75.79921139 | 638559 | 8.937834114 | | | | |
| Scc2-PK Δeco1 Δwpl1 IP_rep1 | 5013750 | 2117407 | 42.23200199 | 1460546 | 29.13081027 | 0.12430746 | 1.270218895 | 1905329 | Fig.S2 |
| Scc2-PK Δeco1 Δwpl1 WCE_rep1 | 6798302 | 4863347 | 71.5376722 | 417007 | 6.133987575 | | | | |
| Scc2-PK wt IP_rep2 | 6392707 | 518654 | 8.113214011 | 5290250 | 82.75445754 | 0.066295358 | 1 | 1500000 | Fig.1, Fig3, Fig.4, Fig.S1, Fig.S2 |
| Scc2-PK wt WCE_rep2 | 7441356 | 4014324 | 53.94613562 | 2714526 | 36.47891594 | | | | |
| Scc2-PK Δeco1 Δwpl1 IP_rep2 | 12267808 | 944791 | 7.70138398 | 10649111 | 86.80532822 | 0.071064364 | 1.071935745 | 1607904 | Fig.1, Fig3, Fig.4, Fig.S1, Fig.S2 |
| Scc2-PK Δeco1 Δwpl1 WCE_rep2 | 13558489 | 7513487 | 55.41537114 | 6018261 | 44.38740187 | | | | |
| Scc2-PK Δwpl1 IP | 4909748 | 989811 | 20.1601182 | 2872937 | 58.51495841 | 0.055980069 | 0.84440405 | 1266607 | Fig.1 |
| Scc2-PK Δwpl1 WCE | 6972689 | 5325310 | 76.37383512 | 865271 | 12.40943057 | | | | |
| Scc2-PK wt_S IP | 2898231 | 1183230 | 40.82593831 | 1305152 | 45.03271133 | 0.027656719 | 0.417174294 | 625762 | Fig.2 |
| Scc2-PK wt_S WCE | 4158198 | 3497417 | 84.10895777 | 106694 | 2.565871082 | | | | |
| Scc2-PK Δeco1 Δwpl1_S IP | 2748831 | 980774 | 35.6796762 | 1340074 | 48.75068711 | 0.026799413 | 0.404242678 | 606365 | Fig.2 |
| Scc2-PK Δeco1 Δwpl1_S WCE | 4141942 | 2990835 | 72.20851958 | 109516 | 2.644073722 | | | | |
| Scc2-PK Pds5-AID_EtOH IP | 4911028 | 2983248 | 60.74589679 | 1239974 | 25.24876665 | 0.061036733 | 1 | 1500000 | Fig.1 |
| Scc2-PK Pds5-AID_EtOH WCE | 5799281 | 5314736 | 91.6447401 | 134833 | 2.324995116 | | | | |
| Scc2-PK Pds5-AID_IAA IP | 6147883 | 340848 | 5.544152353 | 5105647 | 83.04723756 | 0.05513342 | 0.903282615 | 1354924 | Fig.1 |
| Scc2-PK Pds5-AID_IAA WCE | 6550227 | 3082080 | 47.0530258 | 2545358 | 38.85908076 | | | | |
| Scc2-PK wt_Glu IP | 2158161 | 573276 | 19.42885633 | 1584885 | 50.55966631 | 0.180878384 | 1 | 1500000 | Fig.S3 |
| Scc2-PK wt_Glu WCE | 4063358 | 2708800 | 58.52585472 | 1354558 | 28.5462418 | | | | |
| Scc2-PK gal-ub-eco1_Glu IP | 4526737 | 2913293 | 64.35746101 | 1150940 | 25.42537815 | 0.131435076 | 0.726648881 | 1089974 | Fig.1, Fig.S3 |
| Scc2-PK gal-ub-eco1_Glu WCE | 5497945 | 4953896 | 90.10450268 | 257233 | 4.678711773 | | | | |
| Pds5-PK wt IP | 3880013 | 2943690 | 75.86804477 | 254119 | 6.549436819 | 5.37904526 | 1 | 1500000 | Fig.S4 |
| Pds5-PK wt WCE | 4559051 | 2811051 | 61.6586873 | 1305326 | 28.6315288 | | | | |
| Pds5-PK Δwpl1 Δeco1 IP | 4884452 | 3351888 | 68.62362451 | 860322 | 17.61348049 | 3.158911687 | 0.587262522 | 880894 | Fig.S4 |
| Pds5-PK Δwpl1 Δeco1 WCE | 4610114 | 1993369 | 43.23903921 | 1616206 | 35.05783154 | | | | |
| Scc1-PK wt_Glu IP | 5641594 | 3565032 | 63.19192767 | 1477868 | 26.19592973 | 1.945643469 | 1 | 1500000 | Fig.S3 |
| Scc1-PK wt_Glu WCE | 6147371 | 3234123 | 52.6098555 | 2608507 | 42.43288716 | | | | |
| Scc1-PK gal-ub-eco1_Glu IP | 3367198 | 1612993 | 47.90312301 | 1004300 | 29.82598588 | 1.613099911 | 0.829082994 | 1243625 | Fig.S3 |
| Scc1-PK gal-ub-eco1_Glu WCE | 5740738 | 2734643 | 47.63573952 | 2746584 | 47.84374413 | | | | |
| Scc1-PK wt IP_rep1 | 10415274 | 4190427 | 40.23347826 | 5237703 | 50.28867219 | 1.540681638 | 1 | 1500000 | Fig.1, Fig.3, Fig.4, Fig.S1, Fig.S2 |
| Scc1-PK wt WCE_rep1 | 8239307 | 2625194 | 31.86183013 | 5055416 | 61.35729619 | | | | |
| Scc1-PK Δeco1 Δwpl1 IP_rep1 | 18067791 | 8023149 | 44.40581032 | 10033074 | 55.53016415 | 1.064940649 | 0.691213955 | 1036821 | Fig.1,Fig.3, Fig.4, Fig.S1, Fig.S2 |
| Scc1-PK Δeco1 Δwpl1 WCE_rep1 | 21558985 | 9245763 | 42.88589189 | 12312814 | 57.11221563 | | | | |
| Scc1-PK wt IP_rep2 | 15007440 | 6986349 | 46.55256993 | 7939155 | 52.90146088 | 0.893368225 | 1 | 1500000 | Fig.S2, Fig.S8 |
| Scc1-PK wt WCE_rep2 | 19388527 | 9619991 | 49.61692551 | 9766280 | 50.37143874 | | | | |
| Scc1-PK Scc2-AID_G2/M IP | 7532484 | 2497240 | 33.15294131 | 3730839 | 49.52999568 | 0.661045065 | 0.739946919 | 1109921 | Fig.S2, Fig.S8 |
| Scc1-PK Scc2-AID_G2/M WCE | 6215713 | 2631317 | 42.33330915 | 2598666 | 41.80801141 | | | | |
| Scc1-PK Scc2-AID_S IP | 7245745 | 1346198 | 18.57915232 | 4337777 | 59.86654236 | 0.248702003 | 0.278386892 | 417581 | Fig.S8 |
| Scc1-PK Scc2-AID_S WCE | 8151344 | 3881665 | 47.6199385 | 3110682 | 38.16158415 | | | | |
| Scc1-PK Δeco1 Δwpl1 IP_rep2 | 15026713 | 4521746 | 30.09138459 | 10443130 | 69.4971016 | 0.780452492 | 0.873606728 | 1310411 | Fig.S8 |
| Scc1-PK Δeco1 Δwpl1 WCE_rep2 | 25672631 | 9160538 | 35.6821161 | 16511707 | 64.31638035 | | | | |
| Scc1-PK Δeco1 Δwpl1 Scc2-AID_G2/M IP | 15492620 | 3979475 | 25.68626223 | 11429056 | 73.77096966 | 0.600714192 | 0.672414997 | 1008623 | Fig.S8 |
| Scc1-PK Δeco1 Δwpl1 Scc2-AID_G2/M WCE | 19066009 | 6995702 | 36.69201037 | 12069347 | 63.30295449 | | | | |
| Scc1-PK Δeco1 Δwpl1 Scc2-AID_S IP | 8225039 | 1026696 | 12.48256695 | 5921304 | 71.99119664 | 0.173864851 | 0.194617232 | 291926 | Fig.S8 |
| Scc1-PK Δeco1 Δwpl1 Scc2-AID_S WCE | 9318949 | 4337444 | 46.54434744 | 4349318 | 46.67176524 | | | | |
| Rpo21_wt_IP | 3744311 | 3706943 | 99.00200598 | NA | NA | NA | NA | 1500000 | Fig.S2 |
| Rpo21 Δeco1 Δwpl1_IP | 3770581 | 3709096 | 98.36934945 | NA | NA | NA | NA | 1500000 | Fig.4, Fig.S7 |
| Rpo21 Δeco1 Δwpl1_WCE | 4116966 | 3807160 | 92.47489535 | NA | NA | NA | NA | 1500000 | Fig.4, Fig.S7 |

S.cer, Saccaromyces cerevisiae; C.gla, Candida glabrata; OR, occupation ratio; NF, normalization factor

**Supplement Table S4. Micro-C sequencing statistics**

| Sample | Strain | Relative figure | Valid read pairs | Cis | Trans |
|---|---|---|---|---|---|
| wt_rep1 | SKY001 | Fig3, Fig.S9 | 119,690,695 | 84,759,731 | 34,930,964 |
| wt_rep2 | SKY001 | Fig.S9 | 115,657,691 | 75,560,544 | 40,097,147 |
| *Δwpl1 Δeco1*_rep1 | KT110 | Fig3, Fig.S9, Fig.S6, Fig.S7 | 115,443,402 | 80,102,530 | 35,340,872 |
| *Δwpl1 Δeco1*_rep2 | KT110 | Fig.S9 | 107,953,022 | 71,779,182 | 36,173,840 |
| wt *pds5-aid* (EtOH) | SN80 | Fig5 | 86,898,327 | 61,971,243 | 24,927,084 |
| wt *pds5-aid* (IAA) | SN80 | Fig5 | 109,137,056 | 74,506,203 | 34,630,853 |
| *Δwpl1* | KT127 | Fig5 | 124,119,589 | 72,499,681 | 51,619,908 |
| *Δwpl1 Δeco1 scc2-PK* | SN54 | Fig5 | 100,326,853 | 70,685,184 | 29,641,669 |
| wt scc2-aid (EtOH) | SN75 | Fig6, Fig.S8 | 28,410,279 | 18,623,591 | 9,786,688 |
| wt scc2-aid (IAA) | SN75 | Fig6, Fig.S8 | 33,101,775 | 18,319,770 | 14,782,005 |
| *Δwpl1 Δeco1 scc2-aid* (EtOH)_rep1 | SN74 | Fig6, Fig.S9 | 44,163,578 | 28,617,556 | 15,546,022 |
| *Δwpl1 Δeco1 scc2-aid* (EtOH)_rep2 | SN74 | Fig6, Fig.S9 | 83,346,041 | 57,423,809 | 25,922,232 |
| *Δwpl1 Δeco1 scc2-aid* (IAA)_rep1 | SN74 | Fig6, Fig.S8 | 164,153,206 | 77,743,353 | 86,409,853 |
| *Δwpl1 Δeco1 scc2-aid* (IAA)_rep2 | SN74 | Fig6, Fig.S8 | 152,672,879 | 88,652,111 | 64,020,768 |
